# *FUSION*: A web-based application for in-depth exploration of multi-omics data with brightfield histology

**DOI:** 10.1101/2024.07.09.602778

**Authors:** Samuel Border, Ricardo Melo Ferreira, Nicholas Lucarelli, David Manthey, Suhas Kumar, Anindya Paul, Sayat Mimar, Ahmed Naglah, Ying-Hua Cheng, Laura Barisoni, Jessica Ray, Yulia Strekalova, Avi Z. Rosenberg, John E. Tomaszewski, Jeffrey B. Hodgin, HuBMAP consortium, Tarek M. El-Achkar, Sanjay Jain, Michael T. Eadon, Pinaki Sarder

## Abstract

Spatial –OMICS technologies facilitate the interrogation of molecular profiles in the context of the underlying histopathology and tissue microenvironment. Paired analysis of histopathology and molecular data can provide pathologists with otherwise unobtainable insights into biological mechanisms. To connect the disparate molecular and histopathologic features into a single workspace, we developed *FUSION* (**F**unctional **U**nit **S**tate **I**dentificati**ON** in WSIs [Whole Slide Images]), a web-based tool that provides users with a broad array of visualization and analytical tools including deep learning-based algorithms for in-depth interrogation of spatial –OMICS datasets and their associated high-resolution histology images. *FUSION* enables end-to-end analysis of functional tissue units (FTUs), automatically aggregating underlying molecular data to provide a histopathology-based medium for analyzing healthy and altered cell states and driving new discoveries using “pathomic” features. We demonstrate *FUSION* using *10x Visium* spatial transcriptomics (ST) data from both formalin-fixed paraffin embedded (FFPE) and frozen prepared datasets consisting of healthy and diseased tissue. Through several use-cases, we demonstrate how users can identify spatial linkages between quantitative pathomics, qualitative image characteristics, and spatial --omics

## Main

Spatially-resolved, molecular –omics methods are poised to revolutionize medicine both in clinical care and investigative research.^1–3^ These techniques provide investigators with the ability to analyze biological processes down to subcellular resolution within large regions of interest (ROI).^4,5^ When further aligned with histology images, spatial molecular data enrich observations of tissue characteristics and lesions, thereby imparting further insights into localized tissue injury responses. However, the high dimensionality of –omics data represents a barrier toward alignment with histologic findings and paired interpretation. When a pathologist interprets a biopsy specimen, information from large whole slide images (WSI), often consisting of over one billion pixels at 40X magnification, is condensed into a few discrete features which inform treatment and diagnostic decisions. Spatial –omics, by comparison, can contain thousands of deep measurements across the entire tissue area. A similar depth of histologic annotation, wherein every cell or functional tissue unit of an organ is manually characterized at the pixel level, would exceed a pathologist’s clinical capacity. Therefore, paired interpretation of spatial –omics and histology data requires the ability to execute queries of each modality of data, mining heterogeneous information in diverse regions of giga-pixel size. Facilitating this complex integration necessitates a tool, allowing users freedom to simultaneously interact and align multimodal information in a scalable, user-friendly interface, where data dimensionality is tractable and comparable. Such a tool will empower users with automated segmentation of tissue structures using deep learning (DL), quantification of tissue characteristics, dimensionality reduction and statistical analysis of spatial data, and merging both modalities of data in a biologically meaningful fashion. To enhance accessibility and interoperability with other data analysis pipelines, the proposed tool should enable integrated histology and derived data output to share results with the broader research community.

We developed a cloud-based visualization and analysis tool, which combines morphological interpretation of functional tissue units (FTU) at high resolution with genome-wide spatial –omics data. This tool, called Functional Unit State Identification for WSIs (*FUSION*), enables dynamic interaction between users and their data as well as running algorithms with high-computational costs (e.g. GPUs for automated segmentation) utilizing cloud resources (***Fig. 1***). *FUSION* was designed in collaboration with the Human BioMolecular Atlas Program (HuBMAP)^6^ with the goal of aiding users in quantitatively linking spatial –omics datasets to histopathology and determining the distribution of cells and cell states within healthy and diseased tissue. Due to its structural diversity and complex functions, we used the kidney as a prototype organ to demonstrate how *FUSION* can generate easy-to-interpret visualizations of FTUs with their cellular composition. However, this process may also be applied to spatial –omics datasets from all major organs.

**Fig 1.**
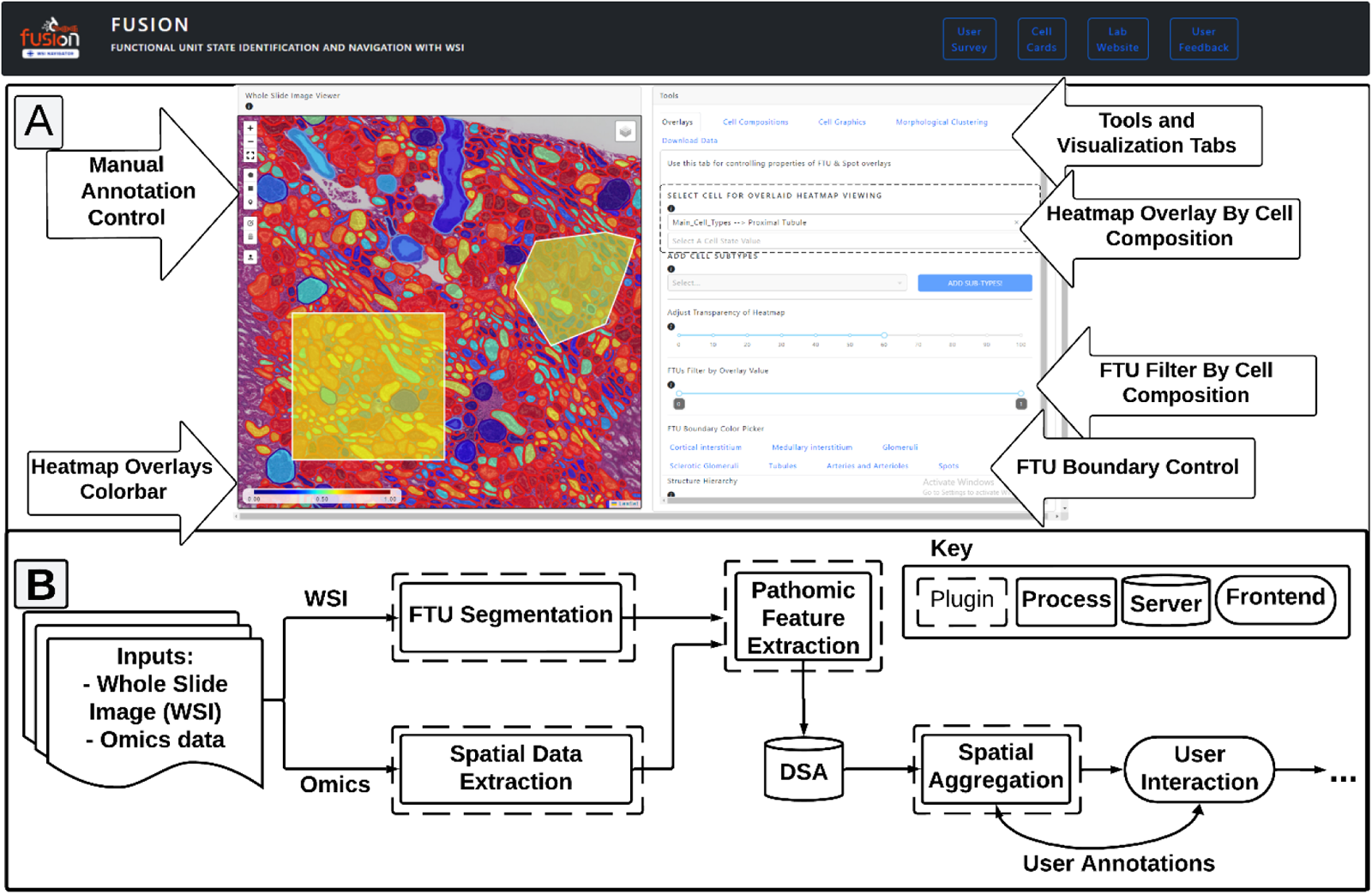
Schematic of FUSION: FUSION offers cloud-based AI solutions unimodal and multimodal analysis of histology images and associated spatial –omics datasets. (A) Frontend visualization of key features of FUSION, demonstrating qualitative and quantitative analysis of spatial tissue maps based on cell composition. (B) Preprocessing flow diagram for separate processing of histological FTUs and spatial –omics data prior to spatial aggregation and user interaction.

## Results

We first discuss a broad system-level overview of *FUSION* (http://fusion.hubmapconsortium.org), and its various functionalities for diverse users, then describe features that are not available in existing cloud tools for large scale image analytics of digital pathology data. We next use paired, multi-modal histology tissue images and associated *10x Visium* spatial transcriptomics (ST) data from kidney tissues of reference (healthy), chronic kidney disease (CKD), and acute kidney injury (AKI) subjects as a vehicle to demonstrate the potential application of *FUSION* in biological research and molecular pathology.

### FUSION

A schematic of *FUSION* is shown in ***Fig. 1***. Broadly, *FUSION* is organized into three pages to facilitate a broad array of biological queries of spatial –omics datasets. These pages represent distinct functions that are crucial to the organization and analysis of large datasets, including data uploading, dataset building, and data visualization. Each page allows users to access plugins which are implemented on the backend through a running instance of Digital Slide Archive (DSA).^7^ On the data uploader page, users can click and drag WSI and associated spatial –omics data files prior to adding slide-level metadata, uploading manual annotations, and/or selecting structures to automatically segment using DL (***Fig. 1A***). The *10x Visium* spatial –omics data contains “spot” (55 µm diameter regions of interest or ROIs) by transcript count matrices, which can be converted into proportions of cell subtypes and their states through a method called cell deconvolution^8^ (***Supp. Doc.***). Currently, *FUSION* hosts *10x Visium* ST sample data from both formalin-fixed & paraffin-embedded (FFPE) (*n* = 12) and frozen (*n* = 44) kidney biopsy tissues and includes reference (healthy), CKD, and AKI cases. End-users can upload their own data or navigate the processed and quality controlled (QC’d) data in *FUSION*. Through the dataset builder page, users can assemble their own datasets based on quantitative, aggregated slide characteristics including structure number, abundance of specific cell types within various structures, and morphometric properties and load those slides into a “visualization session” in the data visualization page.

On the visualization page, *FUSION* enables users to visualize pathomic or cellular feature abundance as a spatial heatmap overlaid on top of the tissue, comparing quantitative cell type and state abundance in various segmented FTUs or manually selected regions (**Supp. Fig. 1 & 2**). Aggregated cell composition data for each FTU is automatically determined when the user navigates to a new region in the histology image. Additionally, users can generate quantitative morphometrics by selecting individual or multiple morphological features or cell types in the “Morphological Clustering” tools tab and generate violin plots (for individual features) and scatter plots (for two or more features). When more than two features are selected, *FUSION* uses uniform manifold approximation projection (UMAP) to reduce the selected features into two dimensions. By clicking on individual points or selecting groups of points in the generated data plots, users can view the FTU associated with that point in the plot as well as their cell type and state abundance.

An interactive illustration panel of various regions in the nephron and the corresponding hierarchical ontology is also presented using the framework developed by the Human Reference Atlas (HRA).^9^ By clicking on different regions within the nephron diagram, users can view a simplified graphic depicting the morphology of cells from that region. This feature increases the utility of this tool for users who are learning more about the cells that contribute to kidney functionality.

*FUSION* complies with *FAIR* (findable, accessible, interoperable, reusable) principles. Our approach focusses on modular engineering and systems-level design for computation, scalability, and expansion to enable future integration of novel spatial data from emerging technologies. Codes and documentation for *FUSION* can be accessed at the GitHub repository: <u>github.com/SarderLab/FUSION</u>

### Multi-modal comparison between disease and reference tissues

In the following sections, we present illustrative use-cases of *FUSION* that integrate histology images and *10x Visium* ST data of kidney tissues from reference, CKD (and diabetic) and AKI patients. These data are analyzed separately using sections from 12 FFPE and 23 frozen section slides. For brevity, we only report analysis of FTU-level features in normal and sclerotic glomeruli, which are defined in the ***Supplemental Document***.

#### Global Comparison of Digital Pathology Image Data

Spatial transcriptomics research yields a large amount of multivariate and bulk measurement data. For decades, pathologists have summarized the histopathologic findings of an entire slide to derive a patient’s prognosis or diagnosis. *FUSION* embraces this clinically useful approach by enabling high-level, efficient aggregation of multivariate features across a whole slide image. The *Dataset Builder* page in *FUSION* facilitates this process in an easy-to-use interface, allowing users to filter datasets based on cell composition and morphometric features recorded across structures within each slide (***Fig. 1A***). The resulting filtered tissue sections allow visualization and plot generation as well as “local” analysis of histological characteristics within individual slides. To demonstrate this feature, we “globally” assessed our first dataset consisting of diabetic (*n*_d_ = 4) and reference (*n*_rc_ = 8) FFPE sections by examining morphometric and cellular features in these datasets (***Fig. 2***). The total number of sclerotic glomeruli and percentage of sclerotic glomeruli across all glomeruli in each biopsy specimen was assessed. Diabetic samples exhibited greater total sclerotic glomeruli (*p* < 0.05) and a higher proportion of sclerotic glomeruli (*p* < 0.05). Using aggregated measurements of cell type composition across different FTUs, *FUSION* also lets users globally assess specific cell types. When comparing diabetic and reference samples, one particular cell type of interest is the podocyte, a unique epithelial cell found in the kidney’s glomerulus that may become effaced or injured in diabetic kidney disease.^10,11^ *FUSION* enables the quantitation of podocyte transcripts in each glomerulus by implementing snRNA-seq deconvolution of spatial transcriptomic data. FUSION was able to identify a reduction in podocyte transcriptional content in sclerotic glomeruli and in non-sclerotic diabetic glomeruli compared to healthy reference tissue (*p* < 0.05).^10,11^ This example illustrates how *FUSION* is able to identify and annotate both pathological and molecular changes in DKD.

#### Structure-Level analysis of Digital Pathology Image Data

At a more granular level, *FUSION* enables the analysis of distributions of derived features within structures, and the examination of the spatial relationships between those structures. This type of query identifies structural “neighborhoods” which are associated with certain cell types or histo-morphological features and can provide insights into pathogenetic mechanisms. Users can select FTU features to analyze, generate figures, and perform statistical analysis on features of biological interests. For a single feature, a violin plot is generated showing the distribution of the selected feature across user-selected labels present in the associated metadata. For two features, a scatter plot is generated to assess linearity between the two selected features. For more than two features, *FUSION* dimensionally reduces the assembled feature vectors into a UMAP which preserves relative sample variation.^12^ Users can then use their mouse to select individual data points or “lasso” groups of points to view the image and cell composition associated with that point, or group of points, on the current plot. This type of visualization allows users to associate the quantitative distribution of features with other qualitative findings in individual FTUs. In addition to hypothesis testing, these features can also result in discovery as the clustering is data driven and agnostic and therefore can reveal unanticipated relationships using morphology and molecular datasets.

**Fig. 2.**
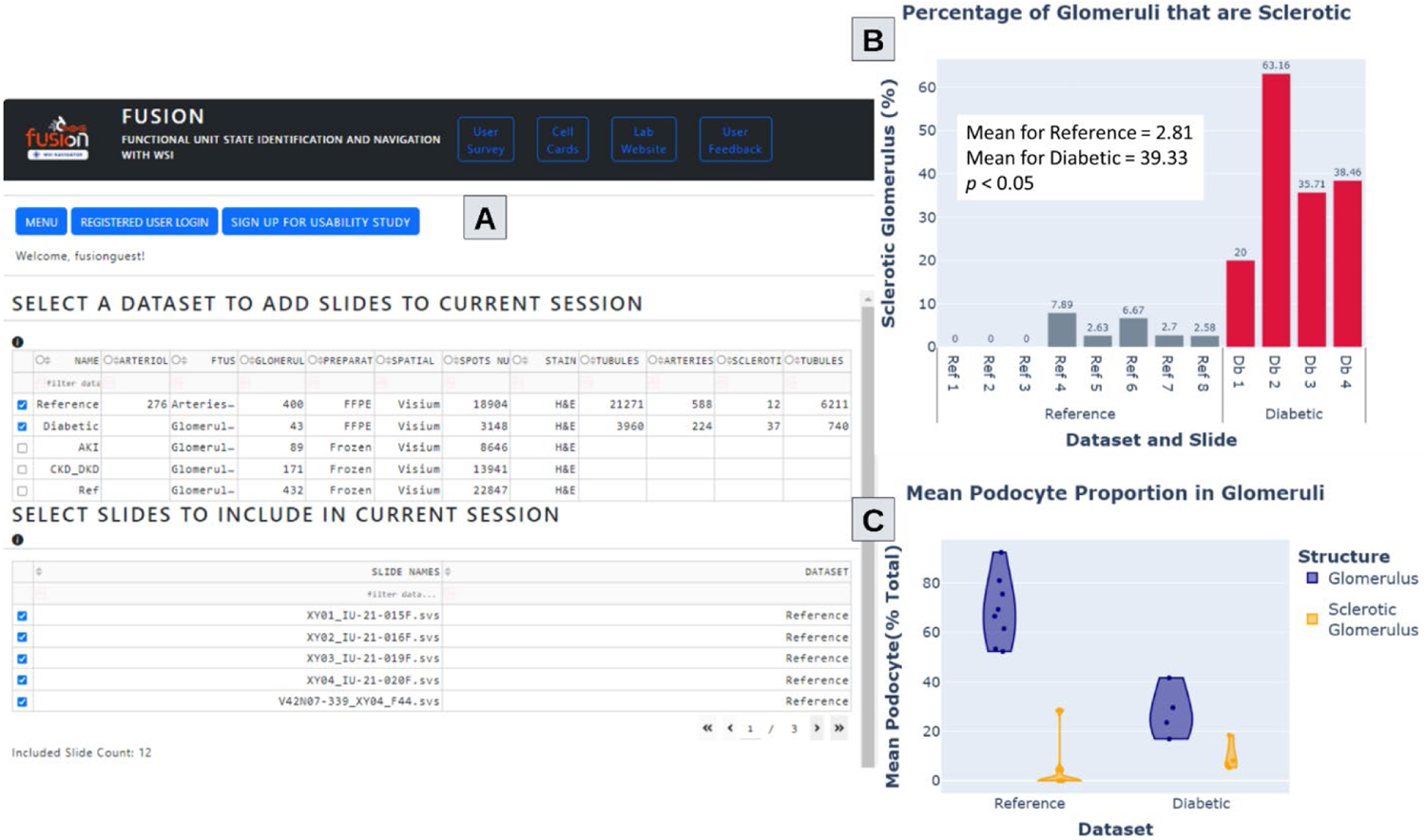
Global comparison of digital pathology image data using Dataset-Builder. (A) Dataset-Builder page in FUSION with all slides from two selected datasets. (B) Percentage of glomeruli that are sclerotic within each slide, separated by dataset (p < 0.05). (C) Proportion of podocytes in sclerotic and normal glomeruli across reference and diabetes samples. Each dot represents a tissue section.

To demonstrate *FUSION’s* ability to succinctly and accurately report on histopathology and molecular features, we examined glomerular hypertrophy in reference and DKD samples. Glomerular hypertrophy, or the inordinate growth in tissue, is a manifestation of DKD resulting from mechanical and chemical stresses. To assess for hypertrophy, we measure the area of eosinophilic tissue within each glomerulus (i.e. histological structure). In glomeruli, these eosinophilic regions comprise cell cytoplasm, mesangial matrix, and basement membranes. As expected, diabetic and sclerotic glomeruli depict higher eosinophilic area (***Fig. 3***),^13^ coinciding with both the physical enlargement of the glomerulus in response to adaptive hyperfiltration, mesangial expansion, and the loss of both Bowman’s space and open capillary lumens in the setting of glomerulosclerosis.^14^

To demonstrate structure-level analysis using multiple morphometric features, we next examine morphological features related to the relative thickness of each sub-compartment. These features are measured using the distance transformation, or the pixel-wise distance from one pixel in a sub-compartment to a pixel of another sub-compartment. Assembled features are first scaled so that each feature has zero mean and unit standard deviation prior to UMAP dimensionality reduction. The resulting scatter plot is then dynamically relabeled in *FUSION* so that the color of each point corresponds to either disease or structure phenotypes (***Fig. 4***). These plots may be used to assess data heterogeneity within respective phenotypes and to make inferences on key features within a particular group.

**Fig. 3.**
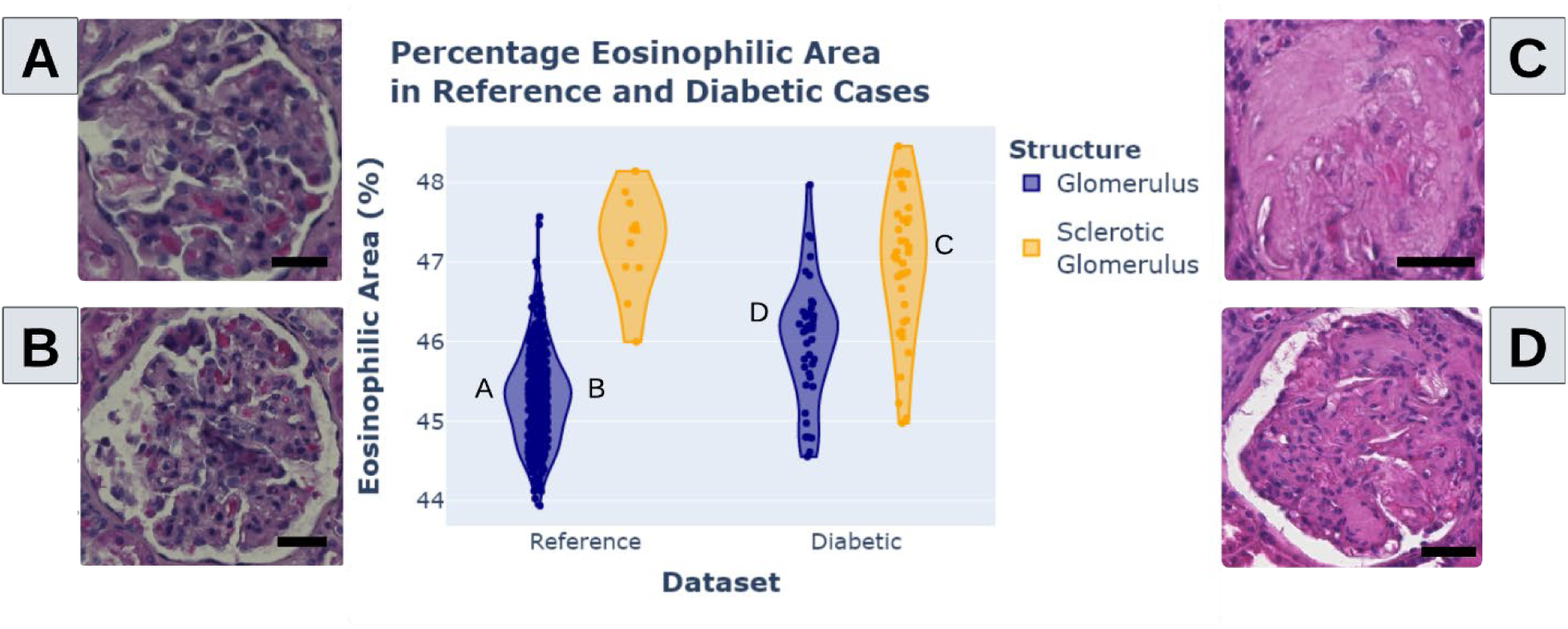
Local comparison of digital histology images using a single feature. As an example, glomerular hypertrophy is compared in diabetes and reference. Percentage eosinophilic area between normal and sclerotic glomeruli from diabetic and reference kidney tissues. (A-D) Example glomeruli selected from the feature distribution where indicated. Dots represent the individual glomeruli. Sclerosis increases eosinophilic area by reducing open capillary lumens and Bowman’’s space. Diabetic kidney disease, in the absence of sclerosis, increases the eosinophilic area through hypertrophy and the expansion of the mesangium. Scalebar indicates 25 µm.

One of the first observations derived from the UMAP plot in ***Fig. 4B-C*** is that the sclerotic glomeruli predominantly group on the left-hand side along with non-sclerotic glomeruli from diabetic cases. To quantify this observed clustering behavior, silhouette scores are automatically calculated in *FUSION* and presented alongside other statistical measures and feature summaries in the *Plot Report* component. For the UMAP containing all glomeruli, silhouette scores of 0.22 and 0.06 are found for the diabetic and reference datasets, respectively, indicating a relatively stronger intra-class grouping of diabetic glomeruli compared to those from reference patients. This observation hints at significant heterogeneity in reference glomeruli, while diabetic glomeruli are concluded to be more homogeneous in this dataset.

*FUSION* enables the assessment of both inter- and intra-subject variability. For example, we observed within the UMAP that the glomeruli from one reference tissue section grouped together more closely. After selecting these points using the lasso tool, the presence of several red (possibly erythrocytes/RBCs) inclusions are identified within glomerular capillary lumens and are likely artifactual related to tissue preparation (***Fig. 4D***). It is common that histopathologic specimens will vary in their preparation, staining character, and image quality. *FUSION* can help to identify artifacts so users can assess whether there is biological relevance or whether artifacts should be excluded or corrected in their analyses.

By selecting other points in the UMAP plot, we can begin to make qualitative conclusions on the relative ordering of glomeruli in the high dimensional scatter (***Fig. 4D***). Specifically, points sampled from right to left on the lower edge of the UMAP plot seem to depict gradual changes from normal glomeruli to fully sclerotic glomeruli. Furthermore, glomeruli towards the upper side of the UMAP plot seem to have a larger overall area. The tool allows discovery of different sub-groups of glomeruli including large sclerotic, small sclerotic, small dense glomeruli, large dense glomeruli, and generally normal glomeruli. Importantly, because FUSION integrates the molecular features (spatial transcriptomics), one can survey molecular expression in these different morphological variants and infer the molecular changes that may underlie these transitions.

The included examples show *FUSION’s* ability to conduct spatial data QC, study data heterogeneity, and find patterns using an interactive approach applicable to disease and reference kidney biopsy sections (***Fig. 2-4***).

#### Local Comparison of Spatial Molecular Data

After using *FUSION* for global and local analysis of image morphometrics, our next focus is on the analysis of spatial molecular data and demonstration of multi-modal data integration.

Among the primary cell types of the glomerulus are podocytes and endothelial cells. As described above, normal glomeruli exhibit a significantly higher proportion of podocyte transcript expression when compared to diabetic glomeruli (*p* < 0.05). Using *FUSION’s* plotting tool, lasso selection tool, and relative cell composition pie charts, a biomedical scientist user can confirm known changes in cell type composition of diabetic glomeruli, wherein the proportion of podocyte to endothelial cell signature shifts (***Fig. 5***). This shift reflects two phenomena: a reduction in podocyte transcript expression related to injury and effacement and an increase in glomerular capillary endothelial cell (EC-GC) transcript expression, potentially resultant from neovascularization in diabetic kidney disease.^15^ Furthermore, a significant difference was found in fibroblast cell fraction between sclerotic and normal glomeruli from both diabetic and reference tissue sections (***Fig. 5B***).

#### Local Comparison Integrating Pathomic Image Features and Spatial Molecular Data

Integrative analysis is achieved in *FUSION* by clustering combined morphometric and molecular features. The first UMAP in ***Fig. 6A*** was generated using only morphometric features from the distance transform and morphological categories. Diabetic samples are loosely grouped toward the left-hand side of the plot, with some glomeruli intermixed with the reference group. Two separate trends are apparent along the spectrum of reference glomeruli (right) to glomeruli (left), indicating two sub-groups of diabetic glomeruli with distinct morphological appearances.

When evaluating glomerulus cell composition features (including non-tubule cell types such as podocytes, aggregate endothelial cells, mesangial cells, immune cells, and fibroblasts) (***Fig. 6B***), we observe heterogeneity in the reference glomeruli resulting in several clusters (upper portion of the plot). Diabetic glomeruli were intermixed, with a subpopulation of glomeruli grouped (bottom right).

**Fig. 4.**
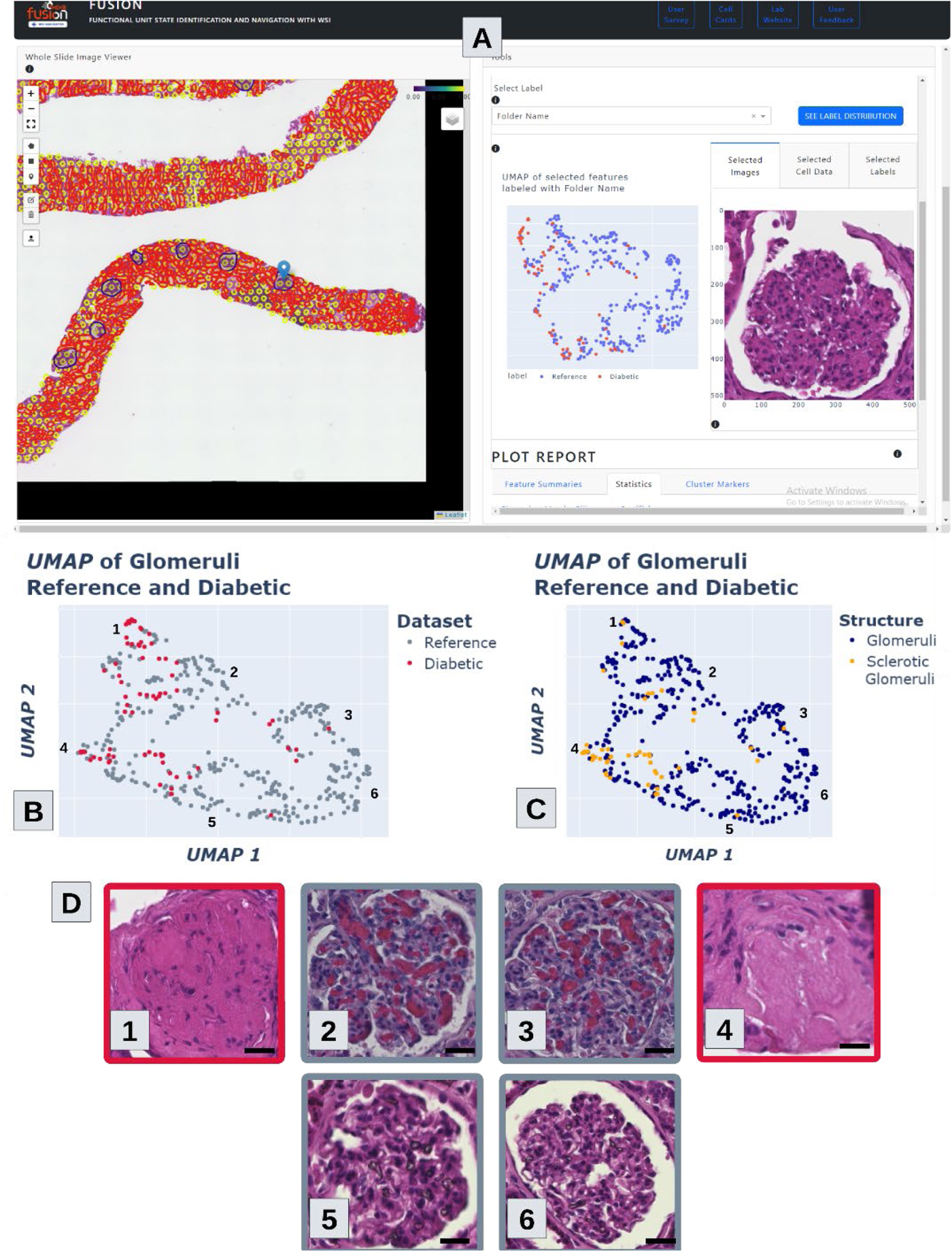
Local comparison of digital histology images using multiple features. (A) Frontend interface of FUSION when selecting multiple features for generating plots. UMAP of glomerular sub-compartment distance transform features labeling (B) whether that glomerulus is from a diabetic or reference slide or (C) whether that glomerulus is a normal glomerulus or a sclerotic glomerulus. (D) Images taken from numbered locations in the above plots. Few different sub-groups of glomeruli are observed. These are large sclerotic, small sclerotic, small dense glomeruli, large dense glomeruli, and generally normal glomeruli. Scalebar indicates 25 µm.

**Fig. 5.**
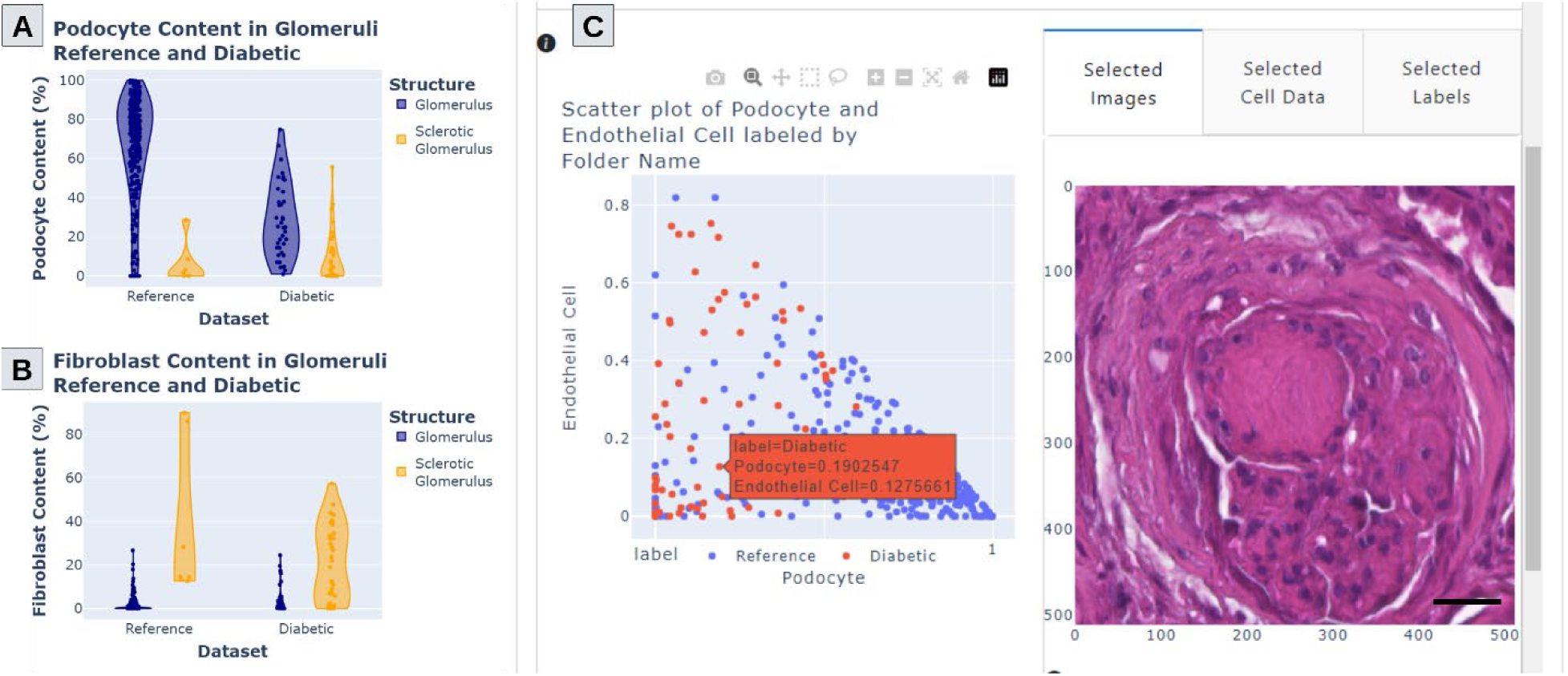
Local comparison of spatial molecular data. Spatial distribution of major cell types in glomerular FTU in our diabetic and reference data. (A) Podocyte proportion in normal and sclerotic glomeruli from reference and diabetic cases. (B) Fibroblast proportion in glomeruli and sclerotic glomeruli from reference and diabetic cases. (C) FUSION frontend example plotting podocyte proportion and endothelial cell proportion with a representative diabetic glomerulus featuring both low podocyte and endothelial cells. Each dot represents a glomerulus. Scalebar indicates 50 µm.

Integrating these two modes of data resulted in the strongest separable clustering of diabetic and reference glomeruli. Diabetic glomeruli were grouped towards the upper left-hand portion of the resulting UMAP plot while reference glomeruli spreading between the middle to lower-right portions with some intermixing of the two groups in between (***Fig. 6C***). By combining morphometrics and cell composition features, silhouette scores for glomeruli in the diabetic group increased from 0.35 with just morphometrics and 0.37 with just cell composition to 0.53. *FUSION* integrates a common analytical technique found in the *Seurat R* package, *Cluster Markers,* to determine key features which separate different groups in dimensionally reduced clusters. This backend implementation offsets the computational costs of this process away from the visualization server, greatly increasing scalability of this process. By selecting the *“Find Cluster Markers”* button, users can send all data that is used to generate a plot to the backend server in order to determine representative features and their adjusted *p*-values. In this example of diabetic and reference glomeruli, the significant discriminative features for glomeruli from the diabetic group include both mean and maximum distance transform values for eosinophilic regions (normalized by object area), increased contents of fibroblasts and immune cells, and nuclear morphology. Specifically, some measures of nuclear aspect ratio indicated a departure from circular nuclei to a more elliptical shape. These features consistently demonstrate a change from a normal glomerulus phenotype to that of a typical diabetic glomerulus, often with associated sclerosis/fibrosis, inflammation, and changes in cellular composition. Glomeruli from the reference group exhibited a higher proportion of podocyte transcript expression and reduced fibroblast transcript expression. To query reproducibility, we repeated the clustering of ***Fig. 6*** using slide identity phenotypes as labels of the respective glomeruli and found the expected cluster patterns (***Supp.*** Fig. 3-5).

**Fig. 6.**
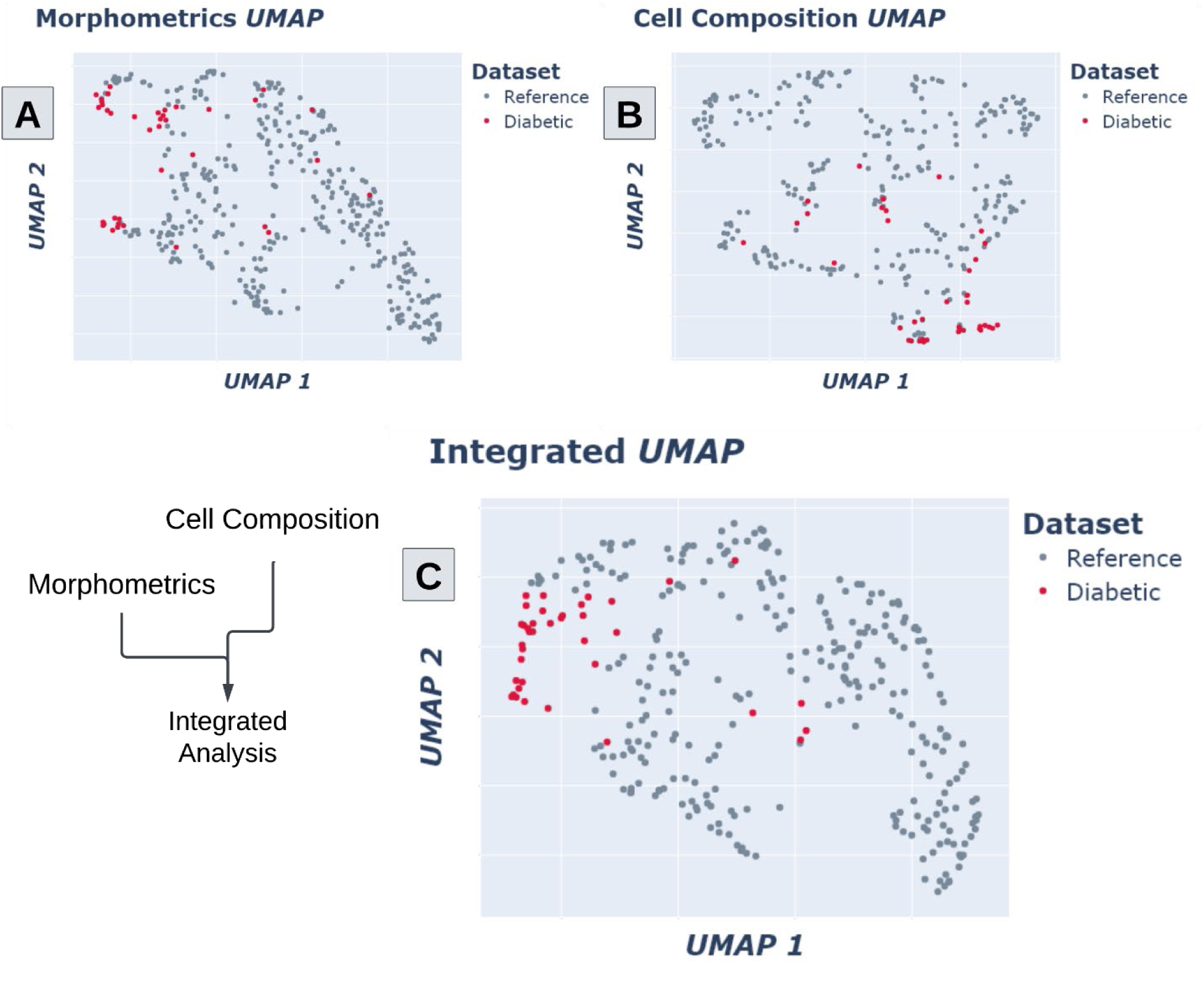
Multimodal data FUSION integrating image feature and cell abundance data derived from spatial –omics data. Glomerular features from both reference and diabetic glomeruli were used to generate UMAP plots. (A) UMAP generated using only morphometric features. (B) UMAP generated using only cell composition features. (C) UMAP generated using a combined set of morphometric and cell composition features. Each dot represents a glomerulus.

#### Spatial Visualization of Immune Cell Infiltration

Thin FFPE sections generally have improved morphology over thick frozen sections. Despite this, *FUSION* has the sensitivity to distinguish some morphologic features in frozen tissue. In the next use case, 23 kidney samples prepared using a frozen tissue section protocol are derived from individuals with AKI (*n*_a_ = 6), CKD (*n*_c_ = 11), and reference (*n*_r_ = 6). The presence and characteristics of inflammation are known to vary greatly between these conditions.^16–19^ AKI, being an acute pathology, often features acute inflammation in the form of infiltrating lymphocytes, but fewer chronic fibrotic changes.^16,17^ CKD, by contrast, shows evidence of continual injury over a longer time, manifested as fibrosis throughout the tubulointersitium in the underlying tissue.^20,21^ The reference tissues may feature some age-related glomerulosclerosis or tubulointerstitial fibrosis, but typically have minimal inflammation.^22,23^

In ***Fig. 7A***, an example with an overlaid heatmap depicts the *10x Visium* spots and glomeruli with greater than 30% immune cell composition. Strong localization of immune cells is seen around a central glomerulus. Fully sclerotic glomeruli in the lower portion of the image exhibit comparatively less immune cell infiltration. This use case illustrates that *FUSION* is able to identify FTUs and regions of inflammation even with the sub-optimal morphology of a frozen section.

**Fig. 7.**
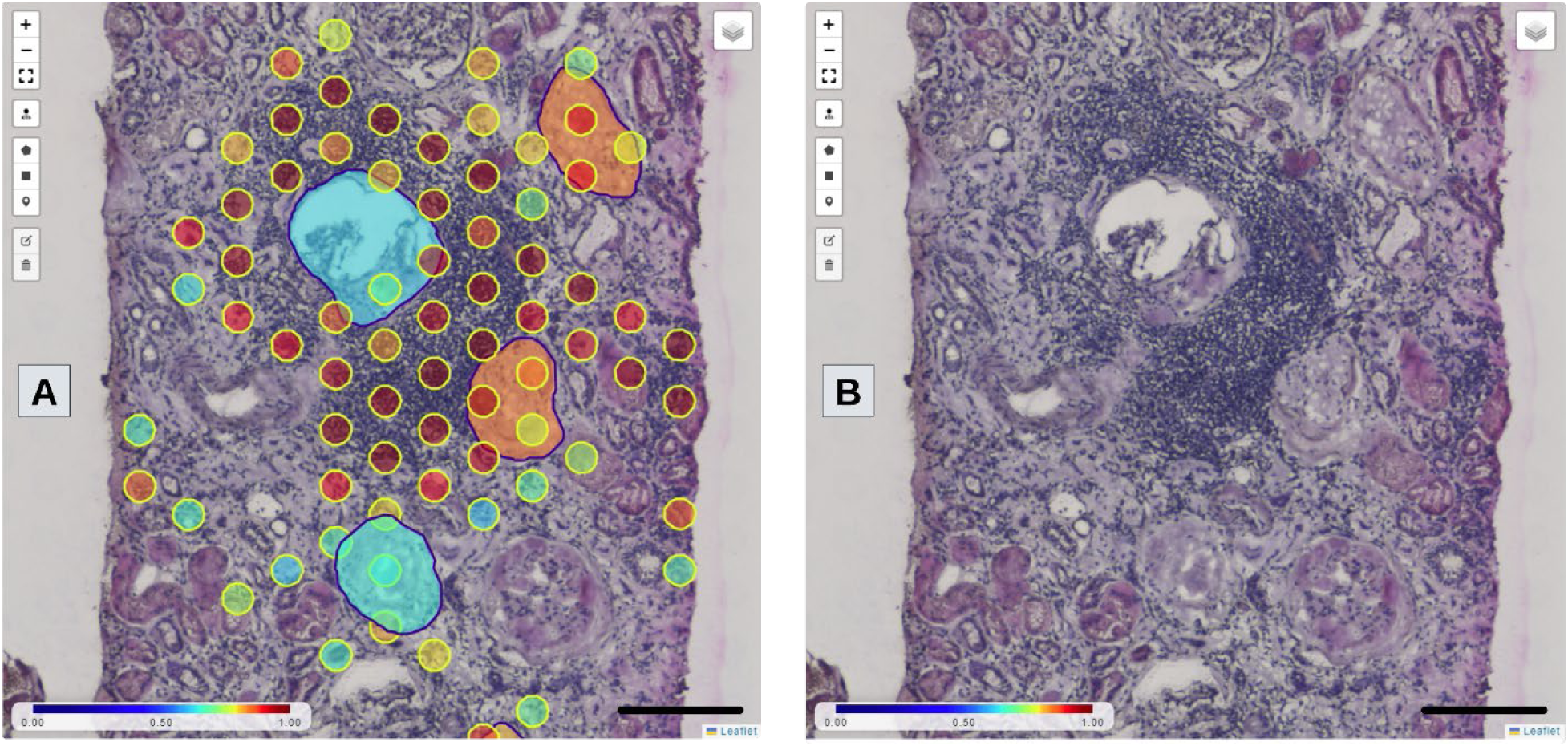
Spatial visualization of immune cell infiltration. (A) Screenshot of a CKD section in FUSION with heatmap set to indicate proportion of immune cells. (B) Same view as in (A) but with annotations turned off for better view of underlying histology. Scalebar indicates 200 µm.

#### Integrating Pathomics and Cell Composition to Study the Impact of Inflammation

Morphological properties can be selected in the clustering tab of *FUSION* along with non-tubule cell types to generate UMAP visualizations (***Fig. 8***). Reference glomeruli clustered distinctly from those of diseased samples (***Fig. 8A***). Cluster markers for these groups revealed statistically significant increased content of immune cells in both AKI and CKD groups compared to the reference glomeruli while the fibroblast content helps to distinguish CKD glomeruli from the AKI glomeruli. *FUSION* enables users to determine the key features driving clustering (***Fig. 8B***). In the *Reference* group, the primary source of heterogeneity between glomeruli stemmed from the different proportions of both podocytes and endothelial cells, both cell types are crucial to normal glomerular function. As expected, the podocyte content of glomeruli in the AKI group (mean = 35.2%) was not substantially lower than that of the reference group (mean = 45.4%). This is expected as changes in podocyte and endothelial cells are often chronic processes, not acute.

In contrast, more immune cell infiltration was found in AKI glomeruli (mean = 16.1%) compared to reference glomeruli (mean = 1.9%). An increased proportion of fibroblast transcripts were found in CKD glomeruli (mean = 10.9%) compared to both AKI (mean = 6.9%) and reference (mean = 3.2%) which aligns with the increased fibrosis in these samples (***Fig. 8C-D***).^16,17^ See in ***Supp.*** Fig. 6 the same UMAPs in ***Fig. 8A-*B** but using slide identity phenotypes as labels of the respective glomeruli.

**Fig. 8.**
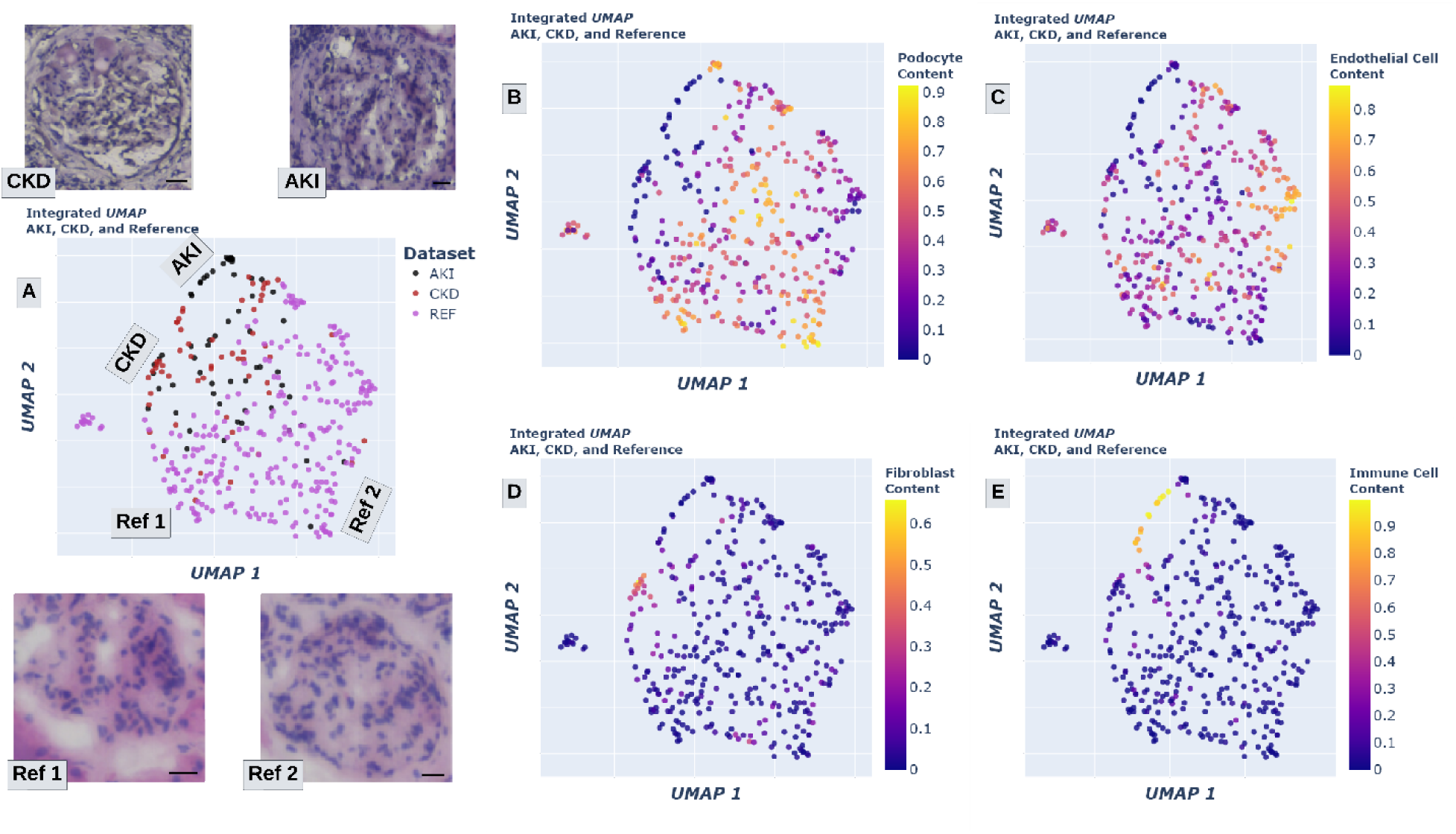
Integrating pathomics and cell composition to study the impact of inflammation. (A) A UMAP plot generated using combined non-tubule cell type composition and morphologic features of glomeruli from AKI, CKD, and reference groups. Select glomeruli pulled out from indicated locations. Each dot represents a glomerulus. (B-E) Same UMAP as in (A) but with the color of each dot corresponding to the content of (B) podocyte, (C) endothelial cell, (D) fibroblast, and (E) immune cell transcripts. Scalebar indicates 25 µm.

## Discussion

We present *FUSION*, a novel visualization, analytic and interactive platform for spatially resolved molecular data that combines histomorphological evaluation with spatial –omics. *FUSION* empowers a wide variety of users to incorporate spatial –omics into their workflows by providing them with user-friendly tools to reveal underlying pathobiological mechanisms. This process serves to increase the accessibility of quantitative computational analyses to a larger user base which normally requires specific skills.

Continuously linking derived data with the source histology image allows for broad associations of quantitative features with established pathological lesions. For a clinical pathologist, this type of interaction enables more direct comparison of segmented structures from one tissue section with those in other sections. When presented with information from many tissue sections, the extensive data visualization methods implemented in *FUSION* provide supportive contextual information which can help clarify ambiguity associated with disease severity grading. This is especially true if longitudinal data are available for those other tissue sections in the dataset. An additional benefit of continuously enabling image and quantitative feature association is that it allows users to validate whether a given image morphometric or textural feature is valuable or robust in distinguishing different types of structures or different diseases. Outlier detection is also possible by identifying data points which correspond to images that contain tissue artifacts such as a fold or tear or presence of red blood cells which can interfere with accurate feature calculation. By enabling users to interact with their data at both the level of a single tissue section as well as for dataset comprising of a set of tissue sections, the efficiency and rigor of biological discovery is enhanced.

One strength of *FUSION* is its extensibility to other modes of spatial –omics. In this work, we present several examples using *10x Visium* spatial data which consists of many ROIs covering the tissue. To effectively translate this data to provide users with structural cell composition, we implemented specialized analytical methods for spatial aggregation of molecular data. These aggregation techniques can be expanded for technologies that provide single-cell resolution such as *10x Xenium*, *CosMx*, and Co-Detection by Indexing (*CODEX*), providing direct insights into cellular composition of complex FTUs.^24–26^ Moreover, our tool is generalizable for any organ systems with spatial and histology data (***Supp.*** Fig. 7), and such data can be accessed through the HuBMAP portal.^27^

It is noteworthy to discuss one limitation of FUSION. Namely, due to the requirement for large, annotated, single-nucleus RNA-sequencing (snRNAseq) datasets, cell deconvolution is currently only available for kidney.^1^ We expect that this deconvolution operation can be expanded for other organ systems with the availability of large scale and comprehensive snRNAseq atlas for other organ systems in the near future.

Through integrations with DSA, *FUSION* enables users to run complex and computationally intense machine learning algorithms on their own data. The *Slicer CLI* plugin framework provides computational researchers with a method to share their pipelines with the community in accordance with FAIR principles.^28^ Further development into the addition and design of new plugins for feature extraction, segmentation, and analysis of spatial –omics data will be the primary aim of future iterations of *FUSION*.

## Author Contributions

SPB conceived the idea of *FUSION,* wrote the code and implemented, and wrote the manuscript. NL assisted in data curation and preprocessing pipelines for spatial –omics data. SK, AP, DM, SM contributed to system infrastructure and implementation of *FUSION*. YS and JR assisted in usability-focused design of different components in *FUSION* for various user groups. AZR, JBH, and LB provided insights from the perspective of research and clinical renal pathologists. JET contributed to the development of data visualization approaches for hand crafted features. Critiqued and edited manuscript. TME contributed to concept development, data generation and manuscript editing. SJ conceptualized the idea of linking histomorphology with multi-omics, provided data, contributed to the idea of different features in *FUSION* and edited the manuscript. MTE, YHC, and RMF generated the *Visium* and *Xenium* data, provided biological interpretation, contributed to feature development within *FUSION*, and edited the manuscript. PS conceived the idea of *FUSION*, helped with study design, coordinated with the study team, mentored SPB, and wrote and edited the manuscript.

## Acknowledgement

We thank the Kidney Precision Medicine Project for providing some of the kidney disease data. Lillian Atchison helped with manuscript editing for final submission. Some of the icons in the cell diagrams are informed and inspired by KPMP.org, Biorender.com, and the Human BioMolecular Atlas Program (HuBMAP).^9^ TME, SJ, MTE, and RMF were supported by U54DK134301. PS’s work was funded by NIH funding from NIDDK - R21 DK 128668 & R01 DK 114485, and from OD OT2 OD033753.

## Methods

### Datasets

**Table 1.**
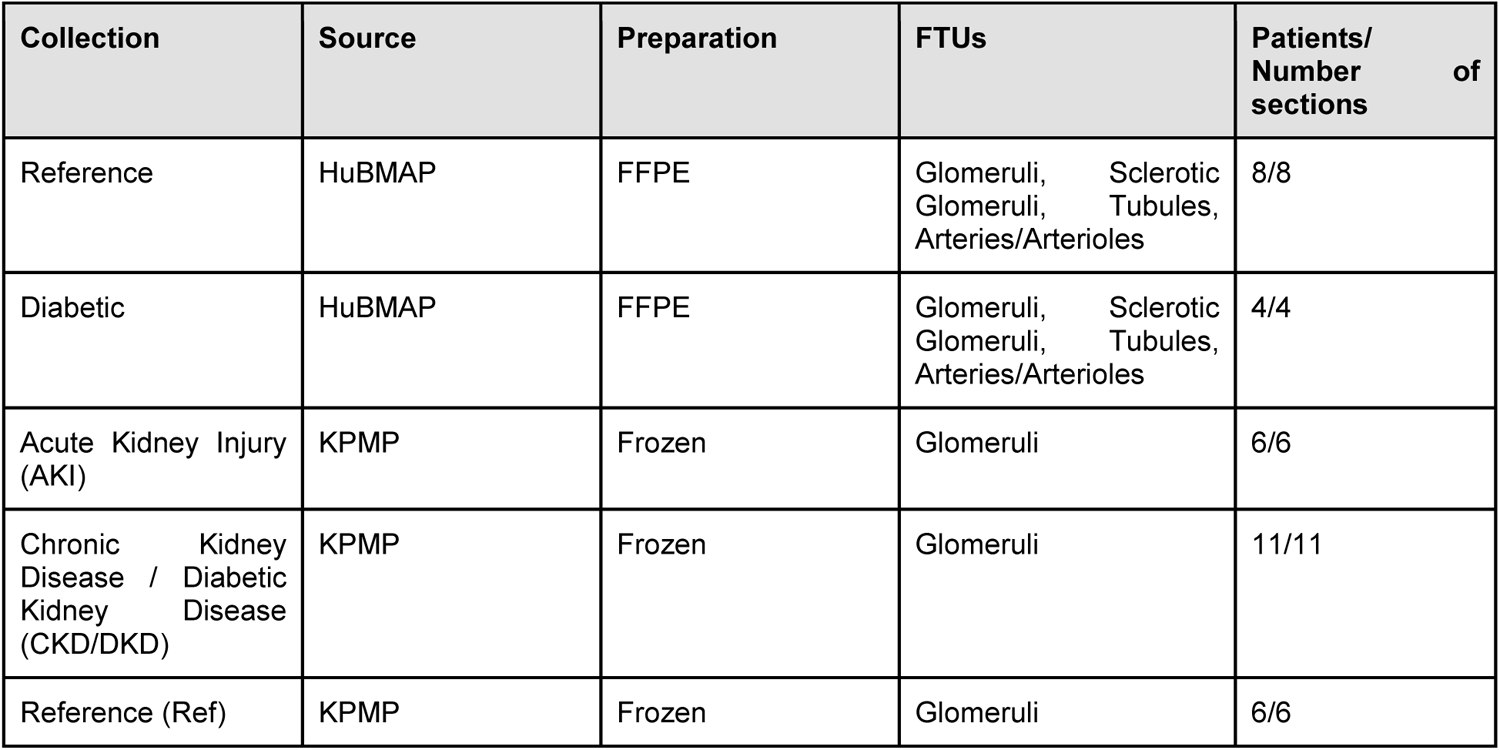
Datasets used in this work. HuBMAP: Human Biomolecular Atlas Program consortium.^29,30^ KPMP: Kidney Precision Medicine Project.^31,32^

#### Input Data Structure

*10x Visium* data used in the development of *FUSION* consisted of two separate types of files as input into the pre-processing steps. The first set of files were the high-resolution scanned WSIs. WSIs varied in file extension depending on the brand name of the scanner that was used or due to downstream post-processing (OME-TIFF). Reading these different types of image files can be accomplished through a number of open-source libraries including *OpenSlide*, *Tiffslide*, and *large-image* which is developed by *Kitware.*^7,33^ These libraries are capable of opening and extracting image regions for a large variety of file formats. *FUSION*, through DSA, uses the *large-image* Python library to read uploaded WSIs, and generate tile sources which are then exposed through an endpoint in the DSA WebAPI.^7^

The second set of files contain transcriptomics read counts and spot center coordinates. These can either be *R* Data Serialization (RDS) files (“.rds” file extension) or “h5ad” files downloaded from the HuBMAP data portal. RDS files contain the *Seurat* object which is output by the *Load10X_Spatial*() command in *Seurat* (v4.0).^34^ Cell subtype proportions for each spot are generated using the *TransferData* function in *Seurat* using a large atlas of single-nucleus RNA-seq (snRNA-seq) data gathered from reference patients as a part of the HuBMAP and KPMP consortia.^9,31^ After this step, the dimensions of the data per-spot are reduced from ∼17k transcriptomic read counts to ∼70 cell subtypes. These cell subtypes are further grouped together according to a smaller set of main cell types of 16 classes, and a varying number of cell states for each main cell type (***Table 2***). This manual data reduction technique was chosen to simplify available visualizations while maintaining the most important information in the pertaining tissue sections. As the spots that are integral to the *10x Visium* Spatial Transcriptomics technique are much larger than a cell (spot diameter = ∼55 µm), each spot is assigned a proportion value for each main cell type that is related to the transcriptomics reads collected at that location.

Spot-level cell type composition data is stored as metadata within the structure annotation text file along with geometry. In *FUSION*, these are formatted following *GeoJSON* convention, where each spot is a *Feature* within a *FeatureCollection* for each slide. Following automated segmentation of select FTUs, cell type composition data is projected from the spots to intersecting FTUs using a weighted sum, where the weight for each spot is equal to the percentage of FTU-area occupied by that spot. Intersection tests, intersection area, and structural area are calculated using the *Shapely* package in Python.

**Table 2.** Breakdown of main cell types, subtypes, and their states. Kidney cell types and states determined using single cell kidney atlas produced by Lake et. al.^1^

| Main Cell Types | Sub-Types | States |
| --- | --- | --- |
| Ascending Thin Limb (ATL) | ATL, dATL | Reference, degenerative |
| Connecting Tubule (CNT) | CNT, CNT-PC, dCNT, cycCNT | Reference, degenerative, cycling |
| Distal Convoluted Tubule (DCT) | DCT, DCT1, DCT2, dDCT, cycDCT | Reference, degenerative, cycling |
| Descending Thin Limb (DTL) | DTL, DTL1, DTL2, DTL3, dDTL3 | Reference, degenerative |
| Endothelial Cell (EC) | EC, EC-GC, EC-AEA, EC-DVR, EC-PTC, dEC-PTC, EC-AVR, dEC, cycEC, EC-LYM | Reference, degenerative, cycling |
| Fibroblast (FIB) | FIB, MYOF, cycMYOF, M-FIB, dM-FIB, dM-FIB, aFIB, dFIB | Reference, cycling, degenerative, adaptive |
| Immune (IMM) | IMM, B, PL, T, NKT, MAST, MAC-M2, cycMNP, MDC, cDC, pDC, ncMON, N | Reference, cycling |
| Intercalated Cell (IC) | IC, C-IC-A, CCD-IC-A, CNT-IC-A, dC-IC-A, OMCD-IC-A, M-IC-A, tPC-IC, IC-B | Reference, degenerative, transitioning |
| Neural-like Cells (NEU) | NEU, SC/NEU | Reference |
| Papillary Tip Epithelial (PapE) | PapE | Reference |
| Principal Cell (PC) | PC, C-PC, CCD-PC, OMCD-PC, M-PC, dOMCD-PC, dM-PC, IMCD, dIMCD | Reference, degenerative |
| Parietal Epithelial Cell (PEC) | PEC | Reference |
| Podocyte (POD) | POD, dPOD | Reference, degenerative |
| Proximal Tubule (PT) | PT, PT-S1, PT-S2, PT-S3, aPT, cycPT, dPT, dPT/DTL | Reference, adaptive, cycling, degenerative |
| Thick Ascending Limb (TAL) | TAL, aTAL1, aTAL2, M-TAL, dM-TAL, C-TAL, dC-TAL, MD | Reference, adaptive, degenerative |
| Vascular Smooth Muscle/ Pericyte (VSM/P) | VSM/P, MC, REN, VSMC, VSMC/P, dVSMC | Reference, degenerative |

### Analytics

#### Cell Deconvolution

Cell deconvolution for *10X Visium* spatial transcriptomic counts is carried out using the *Seurat* package in R. Following normalization and dimensionality reduction using the *SCTransform*, *RunPCA*, and *RunUMAP* functions, identified labels are mapped back to the reference atlas containing sequencing single nucleus information from over 200,000 cells.^1^ Predicted cell subtype proportions are saved back to the RDS file format and used in conjunction with spot centroid locations to derive spot annotation files containing geometry and cell composition data.

#### Automated Segmentation of FTUs

Segmentation of FTUs is performed using one of a few plugin implementations of DL deployed using the *Slicer CLI* Web schema. Each of these models includes PAS-stained kidney sections in their training data. More details on each of the models can be found in ***Table 3***. Our prior work *HistoCloud* outlines plugin creation using DSA.^7,35^

**Table 3.**
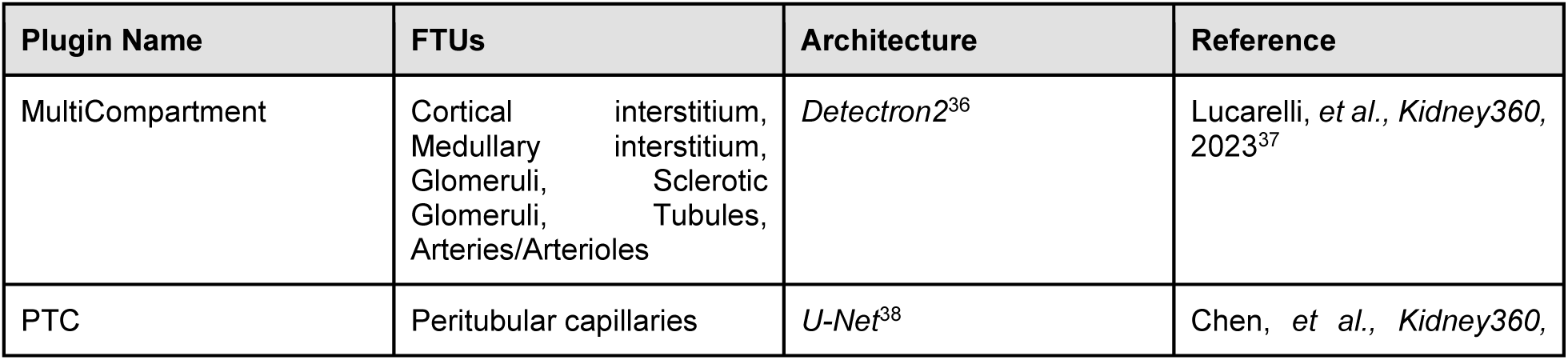
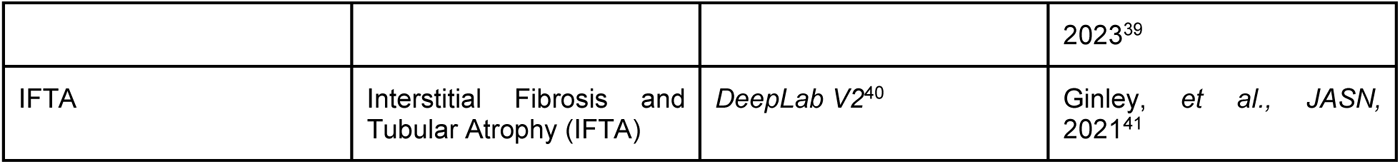
FTU segmentation plugins implemented for *FUSION*.

#### Morphometric Feature Extraction

Morphometrics are calculated for each segmented FTU and added to per-structure metadata alongside cell type composition data. These morphometrics are used in the *Morphological Clustering* tab of FUSION to dynamically cluster structures based on specific morphological characteristics. Morphometrics calculated in this step quantify the size, shape, color and texture of sub-compartments within each segmented FTU. These sub-compartments are segmented using user-defined thresholds during the upload process in an interactive procedure. A full description of these features is provided in ***Table 4***.

**Table 4.** Morphometric feature descriptions. Square brackets ([]) indicate the name of each sub-compartment. The sub-compartment denoting ‘Total’ is the boundary of a structure and all its contents.

| Feature Name | Sub-Compartments | Group | Description |
| --- | --- | --- | --- |
| [ ] Area By Object Area | All | Morphology | Ratio between sub-compartment area and total structural area |
| [ ] Area | All | Morphology | Total area of a given sub-compartment |
| Nuclei Number | Nuclei | Morphology | Number of nuclei contained within a structure |
| Mean Aspect Ratio Nuclei | Nuclei | Morphology | Mean aspect ratio (major axis length divided by minor axis length) across all nuclei within a structure |
| Standard Deviation Aspect Ratio Nuclei | Nuclei | Morphology | Standard deviation of aspect ratio (major axis length divided by minor axis length) across all nuclei within a structure |
| Mean Nuclear Area | Nuclei | Morphology | Mean area of all nuclei within a structure |
| Total Object Area | Total | Morphology | Total area of structure measured in number of pixels |
| Total Object Perimeter | Total | Morphology | Total perimeter of structure boundary measured in number of pixels |
| Total Object Aspect Ratio | Total | Morphology | Aspect ratio (major axis length divided by minor axis length) for a structure |
| Major Axis Length | Total | Morphology | Length of longest axis of ellipse with same normalized second central moments as structure |
| Minor Axis Length | Total | Morphology | Length of shortest axis of ellipse with same normalized second central moments as structure |
| Contrast [ ] | All | Texture | Measures intensity contrast between a pixel and its neighbor over the whole sub-compartment. 0 for 'constant' (An image consisting of one color with no variation). |
| Homogeneity [ ] | All | Texture | Measures the closeness of the distribution of elements in the Gray-Level Co-occurrence Matrix (GLCM). 1 for a diagonal GLCM (constant, single color image) |
| Correlation [ ] | All | Texture | Measures how correlated a pixel is to its neighbor over the whole sub-compartment. 1 or -1 for a perfectly correlated image and a perfectly negative correlated image, NaN for a constant image (constant, single color image). |
| Energy [ ] | All | Texture | Sum of the square elements in the Gray-Level Co-occurrence Matrix (GLCM). 1 for a constant, single color image. |
| Mean Red [ ] | All | Color | Mean of red values (first color channel in a RGB image) across a sub-compartment. |
| Mean Green [ ] | All | Color | Mean of green values (second color channel in a RGB image) across a sub-compartment. |
| Mean Blue [ ] | All | Color | Mean of blue values (third color channel in a RGB image) across a sub-compartment. |
| Standard Deviation Red [ ] | All | Color | Standard deviation of red values across a sub-compartment. |
| Standard Deviation Green [ ] | All | Color | Standard deviation of green values across a sub-compartment. |
| Standard Deviation Blue [ ] | All | Color | Standard deviation of blue values across a sub-compartment. |
| Sum Distance Transform By Object Area [ ] | All | Distance Transform | Sum of the distance transform values divided by total object area for a sub-compartment |
| Sum Distance Transform [ ] | All | Distance Transform | Sum of the distance transform values for a sub-compartment. |
| Sum Distance Transform by [ ] Area | All | Distance Transform | Sum of distance transform values for a sub-compartment divided by total area of the same sub-compartment. |
| Mean Distance Transform By Object Area [ ] | All | Distance Transform | Mean of distance transform values for a sub-compartment divided by total object area. |
| Mean Distance Transform By [ ] Area | All | Distance Transform | Mean of distance transform values for a sub-compartment divided by total area of the same sub-compartment. |
| Mean Distance Transform [ ] | All | Distance Transform | Mean distance transform values for a sub-compartment. |
| Max Distance Transform By Object Area [ ] | All | Distance Transform | Maximum distance transform value for a sub-compartment divided by total object area. |
| Max Distance Transform By [ ] Area | All | Distance Transform | Maximum distance transform value for a sub-compartment divided by the total area of the same sub-compartment. |
| Max Distance Transform [ ] | All | Distance Transform | Maximum distance transform value for a sub-compartment. |

### Frontend

*FUSION* is built using *Dash*, a web-based visualization framework for Python applications.^42^ Components within *FUSION* are derived from either the core components library or several open-source community components including *Dash-Leaflet* (for large image visualization), *Dash-Bootstrap-Components* (for grid-style layouts, icons, and organization), *Dash-Extensions* (for integrating *JavaScript* functions), among others.

The visualization page (http://fusion.hubmapconsortium/vis), contains the majority of interactive components in *FUSION*. These are divided amongst WSI visualization and overlays, interactive plotting, cell graphics and hierarchy viewing, and data downloading. The WSI Viewer card contains the WSI, FTU/spot annotations and their associated overlays, and custom ROIs. Integrated *JavaScript* functions are used to control boundary color, fill color, transparency, and filtration of overlaid FTU and spot annotations according to user inputs. Annotations and image tiles are hosted in DSA and accessed through the annotations and item tiles WebAPI endpoints. By logging in through *FUSION*, users can access their personal uploaded images. Although annotation correction is not implemented for FTUs in *FUSION*, users may correct predicted FTU boundaries using *HistomicsUI*, which is accessed through DSA at the host address. Combined morphometrics and cell composition plotting is available in the *Morphological Clustering* tab. In this tab, users can select features, labels, and filters to generate an interactive plot of data in their current dataset. For single features, violin plots of the raw data are generated. Selecting two or more features results in a two-dimensional scatter plot containing either raw data (for two features) or UMAP coordinates for dimensionally reduced data. Users can select one or multiple points to view the FTU or spot with that feature value. By selecting multiple points, an animation is generated that allows the user to scroll through several images or FTUs at a given location at a time (maximum 100 points, for efficiency) as well as view their mean cell composition. This visualization allows users to make broader connections for qualitative observations to the quantitative morphometric or –OMICS values present in a given FTU. Users can also download the cell composition data for FTUs which are in the current view in either Comma Separated Values (CSV) or Excel (XLSX) format. These files will list each FTU in the current view and have a column corresponding to a particular cell.

From the dataset builder page (http://fusion.hubmapconsortium/dataset-builder), users can manually select which datasets they would like to examine in their current visualization session. We provide 56 pre-processed samples from both FFPE and frozen preparations (***Table 1***) from both healthy, reference and disease groups. After selecting one or more datasets, users can make preliminary visualizations of aggregated metadata including FTU-level cell type composition statistics, counts of each FTU, and other features of interest. This feature can help inform users whether there are certain slides they would like to remove from their current selection. Once a user is satisfied with their collection of slides, they can then click the ‘*Go To Visualization*’ button to be redirected to the visualization page where their selected slides can be viewed, and their clustering features accessed.

Pre-processing workflows are accessed through the data upload page (http://fusion.hubmapconsortium/dataset-uploader). Here, users can select what kind of data they would like to upload, upload the required files, select FTUs to segment, select parameters to use to segment sub-compartments within segmented FTUs, and calculate which morphometric features they would like to calculate. All of which is accomplished using simple user-interface components with descriptive instructions throughout.

### Backend

*FUSION* works in concert with a running instance of DSA.^7^ DSA is a containerized cloud platform which provides users with a secure, flexible, and scalable interface for the analysis of WSIs. Included in DSA is a rich RESTful API (Representational State Transfer Application Programming Interface) and Girder for data management operations. DSA (specifically the Web API) is used by *FUSION* to store pre-processed and newly uploaded data, run *Slicer CLI* Web plugins (specially formatted Docker images), access WSI tiles for large image visualization at multiple levels of resolution, and user registration. Using the *Girder Client* Python library as well as the *requests* Python library, requests can be made to the DSA server from an external host. This means that the deployment of *FUSION* and its corresponding DSA instance can have separate resource allocations/specifications which fit what is expected of frontend and backend servers. For example, WSI segmentation jobs are accelerated using graphics processing units (GPUs), and cell deconvolution requires a significantly higher amount of random-access memory (RAM) compared to rendering visualization components on the frontend.

### Data, Code, Tutorial, & Promotional Video

Codes for *FUSION* may be accessed at the *GitHub* repository: github.com/SarderLab/FUSION. Data used in this project may be accessed through the data portals of participating consortia (portal.hubmapconsortium.org). *FUSION* can be accessed via http://fusion.hubmapconsortium.org/. A promotional video on FUSION is available at https://www.youtube.com/watch?v=d1tHayLENEE

## Supplemental Document

### Glomerulus: Normal and Sclerotic

Glomeruli are the primary filtration units of the kidney, consisting of a bundle of capillaries with fenestrated endothelial cells and specialized, terminally differentiated epithelial cells (podocytes) suspended in a matrix created by mesangial cells together referred to as the glomerular tuft.^43–45^ In addition to the glomerular tuft, healthy glomeruli also feature a large empty space (Bowman’s space) which contains fluid prior to drainage into the lumen of nearby proximal tubules.^46^ Due to the specialized nature of these cells and capillaries, glomeruli are especially sensitive to increases in blood pressure (hypertension) as well as the production of advanced glycation end-products (AGEs) which are both typically present in cases of diabetes mellitus.^47^ As chronic injuries accumulate, capillary loops, cross-sectional views of capillaries in the glomerular tuft, are closed off and overtaken by expanded mesangial matrix and fibrotic scars.^13^ The resulting glomerulus has no ability to filter blood and is rendered inert, becoming known as a sclerotic glomerulus (or globally sclerotic glomerulus). Sclerotic glomeruli are not exclusively a diabetic lesion; glomeruli can also become sclerotic over the course of normal aging.^22,23^

**Supp. Fig. 1.**
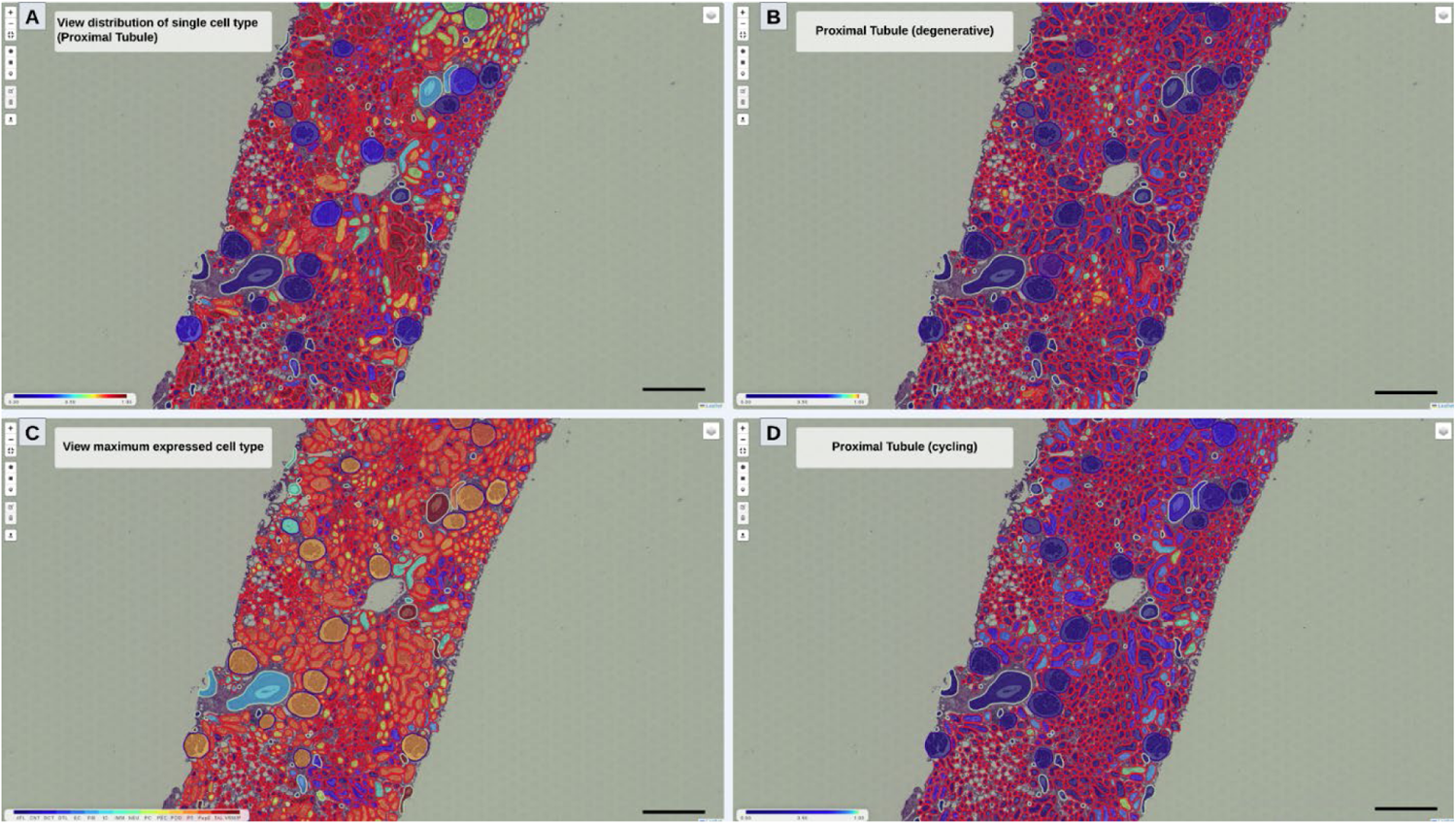
Visualization of cell type and state distribution spatially with respect to FTUs. Overlaid color corresponding to (**A**) proportion of proximal tubule cell type, (**B**) proportion of proximal tubule with the degenerative cell state, (**C**) highest proportion cell type per structure, and (**D**) proportion of proximal tubule with the cycling cell state. Scalebar indicates 500 µm

### Overlaid Visualizations on Histology

Annotations in histology images have the potential to contain a diverse amount of derived data. A primary aim of digital pathology is to quantify underlying patterns in histological structures to increase the objectivity of pathological assessment. One approach is to use a combination of classical and modern image analysis methods to calculate hand-engineered morphometric features for annotated structures. These morphometric features can encompass morphologic quantifications such as size and shape but can also include color, texture, and relative positioning of structures and their sub-compartments. FUSION enables users to calculate a wide range of morphometrics as a part of its preprocessing procedure. This data is combined with spatially aggregated –omics data for correlative analysis of histologic phenotype and gene expression. One way to view spatial distributions of specific cell types and morphological properties is through a heatmap overlaid on the histology WSI. This visualization also allows for users to determine relative similarity between structures as well as which structures are the most different from each other based on a specific value.

Upon loading a new slide in FUSION, the properties which can be used to generate overlaid heatmaps are automatically determined and added to a searchable dropdown menu. These properties include standalone structural metadata properties as well as sub-properties (e.g., Main Cell Types → Podocyte). Selecting one of these properties prompts FUSION to collect the value for that property from all structures where that property is present in their metadata. If this is a numeric property (e.g., proportion of endothelial cells), these values are then arranged from smallest to largest and assigned a color from dark blue (smallest) to dark red (highest) according to the “JET” colormap. For categorical properties (e.g., maximally present cell type), each label is assigned its own color alphabetically with a categorical colorbar indicating each color and its label. In this section, we will present two possible use-cases of overlaid heatmaps and describe how they facilitate efficient deduction.

#### Sub-Categorization of FTUs by Cell Composition

This first application of overlaid heatmaps involves sub-categorization of different FTUs by cell composition. This task is important because it allows users to approximate the functional role of various structures within their image. In spatial transcriptomics, cell types can be defined by the expression of one or more transcripts. When two or more transcripts define a cell type, such as those derived from a single cell RNA-seq cell cluster, these expression signatures can be summarized as a single vector, sometimes called a transfer score. These transfer scores can then be used to determine which cell types are heavily localized along sites of injury or in various regions in the tissue. This information can also be used to establish FTU neighborhoods with shared cell type or cell state proportion within a small area.

After selecting a cell type from a dropdown menu, users can use the FTU filter slider to change the acceptable value range for FTUs. This slider has both a minimum and a maximum value which can be adjusted in response to queries like, “*Show me all structures with a Proximal Tubule cell proportion greater than 0.5.*” Using this filter helps remove unwanted structures from the current slide and allows users to focus more on their desired structures of interest.

In the examples shown in ***Supp.*** Fig. 1, the proportion of proximal tubule is selected first for overlaid viewing followed by the highest proportion cell type. In the proximal tubule view, the majority of tubules in this slide represent the proximal tubule cell type with a smaller proportion of tubules coming from different portions of the nephron. As expected, structures such as glomeruli and arterioles exhibit minimal proximal tubule content. In the maximum expressed cell type view, the tubules that do not contain a large amount of proximal tubule cells seem to contain more thick ascending limb cells. Interspersed throughout the slide, we also see distal convoluted tubules grouped together. Another observation that can be made is that most of the glomeruli have the podocyte as the cell type present in highest proportion. For this particular WSI, a healthy reference sample, the difference in glomerular cell composition is not very large. However, decreasing podocyte fraction is an important factor in glomeruli of certain diseased groups.

#### Cell State Comparison between FTUs

Another application of overlaid heatmaps is to combine multiple FTU properties into one visualization. We demonstrate this functionality using cell state proportions. In the context of kidney, cell states are defined from a large single nucleus RNA-seq (snRNA-seq) dataset consisting of more than 200K kidney cells.^12^ Intra-cluster variation observed for different cell types are further examined for relative presence of specific markers which inform researchers on more specific functions or reactions of those cells to certain stimuli. Possible cell states in kidney data include reference (health), degenerative, adaptive, and transitioning.^12^ Cell state composition is stored for each FTU as a nested sub-property wherein each “*Main Cell Type*” contains a varying set of possible cell states and a value for each state. This can be interpreted as the proportion of each cell type made up by each cell state. A glomerulus can therefore contain, for example, 75% podocyte with 90% of that coming from the reference (healthy) cell state and the remaining 10% belonging to an injury cell state (e.g., degenerative). Combining these two properties is accomplished by multiplying the total percentage of each cell type by the selected cell state. For the earlier example, the glomerulus consists of 67.5% reference podocytes and 7.5% degenerative podocytes. This method of combination is particularly useful for comparison in cell states between FTUs in a WSI, as we are interested in the total content of a particular cell state, and therefore, need to know both the total proportion of a given cell and its composite cell states.

In ***Supp.*** Fig. 1, localization of both degenerative and cycling proximal tubules are observed in various sub-regions throughout the same region of tissue. This observation can be especially useful in identifying localized patterns of injury, stress, or repair mechanisms. Combined overlay visualizations can also be filtered using the same FTU filter slider described in the previous section.

### Defining Custom ROIs for Spatial –omics Aggregation

A key form of interaction with data favored by the pathology community is drawing on or annotating regions or structures of interest in WSIs. These annotations are used for counting, highlighting, sharing findings with collaborators, and secondary analyses. When dealing with spatial –omics data, researchers are also interested in underlying molecular data in regions which are annotated. Therefore, dynamic aggregation of cell composition data is a necessity for spatial –omics interfaces.

In FUSION, aggregation of spatial molecular data is included both as a pre-processing step for automatically segmented FTUs as well as being incorporated into a variety of manual annotation tools (***Supp.*** Fig. 2). Annotation shapes include freeform polygons, rectangles, and markers which are located on the left-hand side of the WSI viewer component. Each of these tools triggers an aggregation function for structures which are enclosed within the annotation boundaries or intersecting with the marker. Freeform polygons and rectangles are used for annotating whole regions of tissue, including regions outside of existing FTU boundaries. This process enables users to collect information for loosely defined areas such as regions of interstitial fibrosis and tubular atrophy (IFTA). After making a manual annotation with either of these tools, the same visualization properties are available as those for FTUs including overlaid heatmap colors and cell composition plotting. Since these manual ROI cell composition charts are static (independent of position on WSI), users can readily compare the cellular makeup of their manually annotated regions with different regions of tissue or FTU groups. Alternatively, users can use the marker tool to select FTUs to group together from different regions in the current WSI. This process ensures that the aggregated cell composition values correspond only to the selected FTUs of interest. FTUs can also be marked automatically from selected data points in the morphological clustering tab that are in the current slide.

**Supp. Fig. 2.**
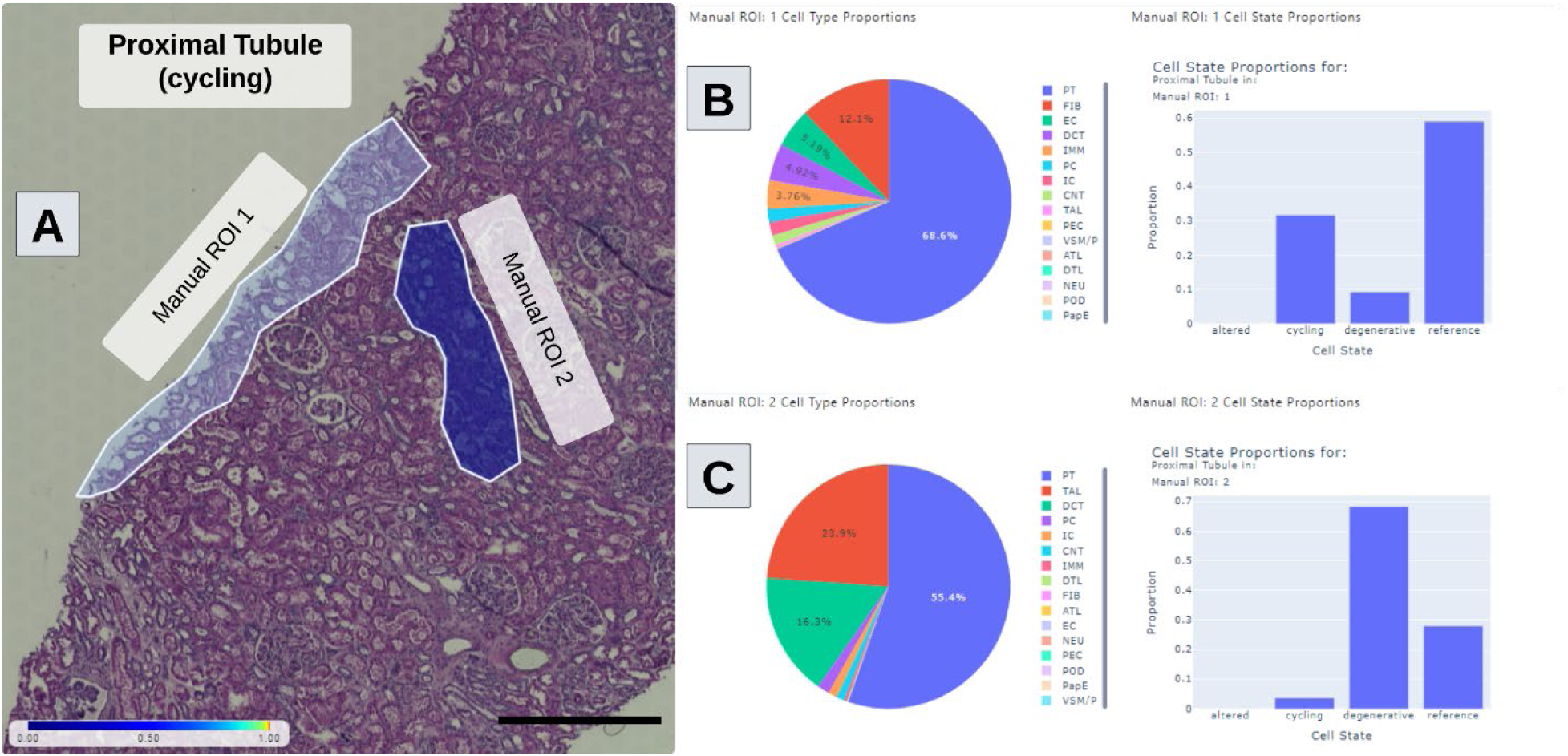
Visualization of cell type and state distribution spatially with respect to arbitrary annotations. (**A**) Two manually annotated regions of interest with overlaid color corresponding to proportion of degenerative proximal tubule. (**B**) Cell composition pie charts for each FTU where each tab indicates the FTU name and number contained within the current viewport. Manual ROI 1 here contains a higher proportion of cycling proximal tubule. (**C**) Cell composition pie chart for Manual ROI 2 indicating a much higher proportion of degenerative proximal tubule cell type than Manual ROI 1. Scalebar indicates 200 µm.

Once a user has manual annotations on a slide, they can also download image and molecular data for selected regions for secondary analysis in the “*Download Data*” tab. Annotations can be downloaded in three different formats for visualization on other platforms such as *Aperio ImageScope*, *QuPath*, and *HistomicsUI.*^19–21^

**Supp. Fig. S3.**
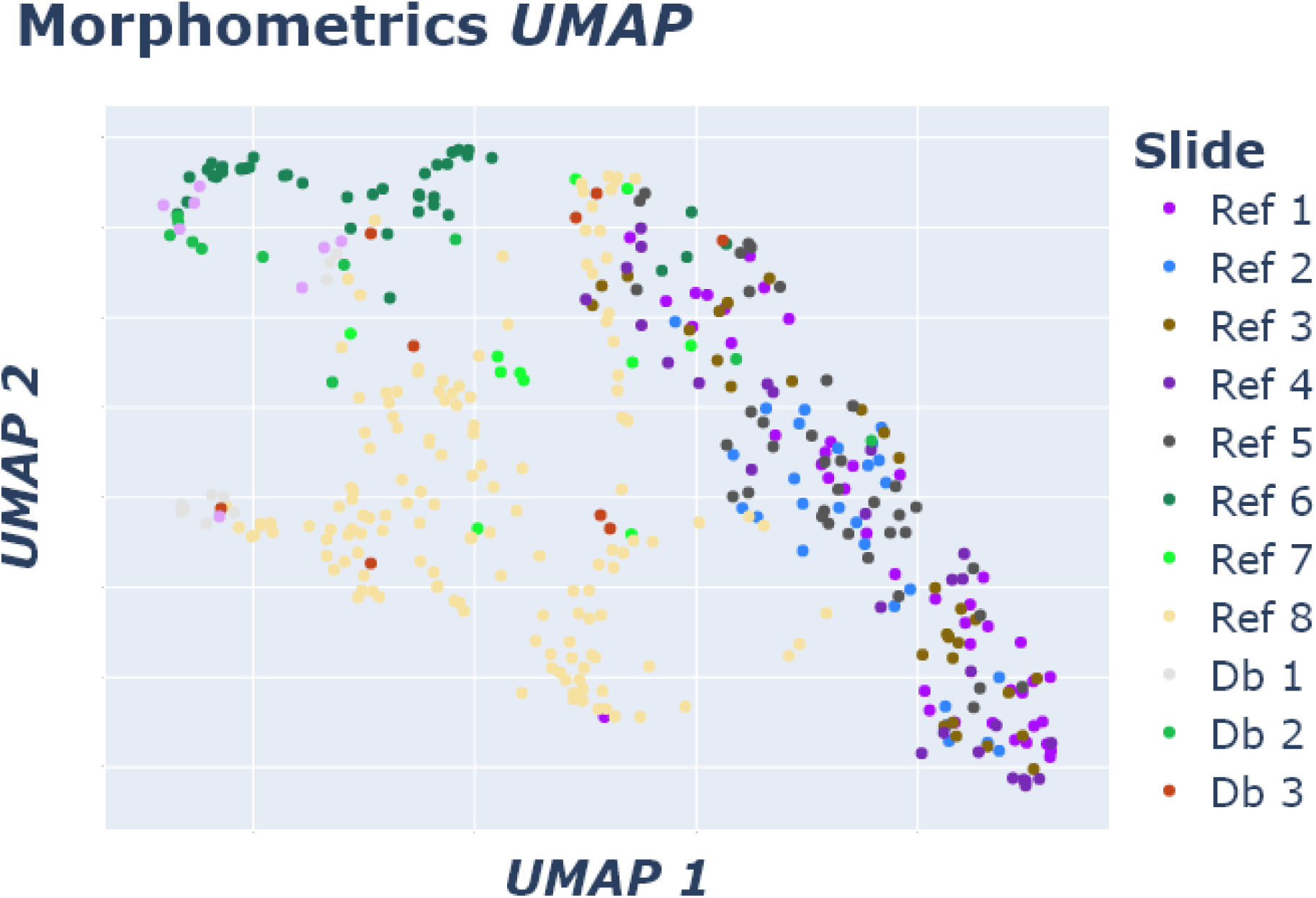
UMAP of morphometric features from reference and diabetic slides. Each dot indicates a glomerulus.

**Supp. Fig. S4.**
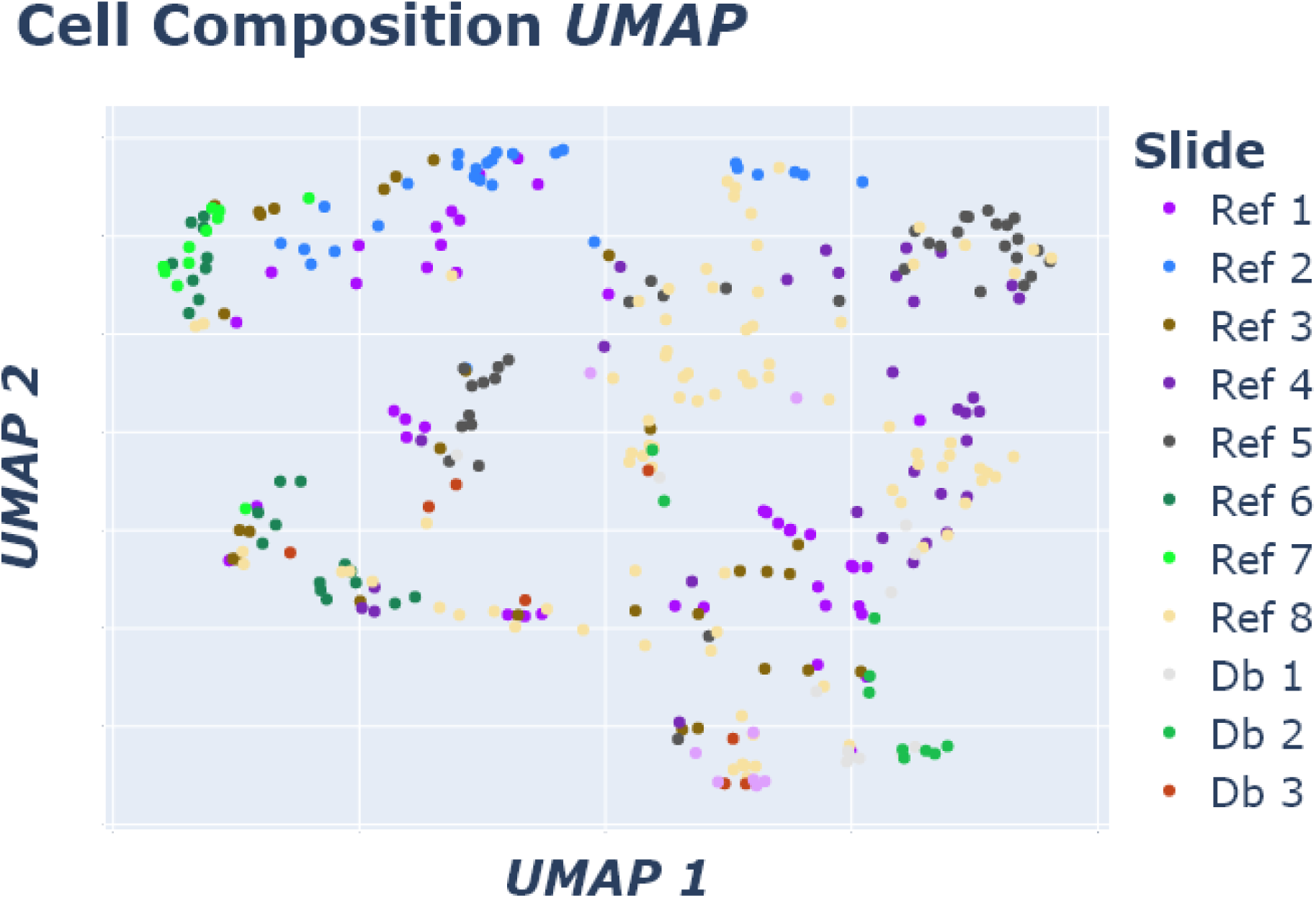
UMAP of cell composition features from reference and diabetic slides. Each dot indicates a glomerulus.

**Supp. Fig. S5.**
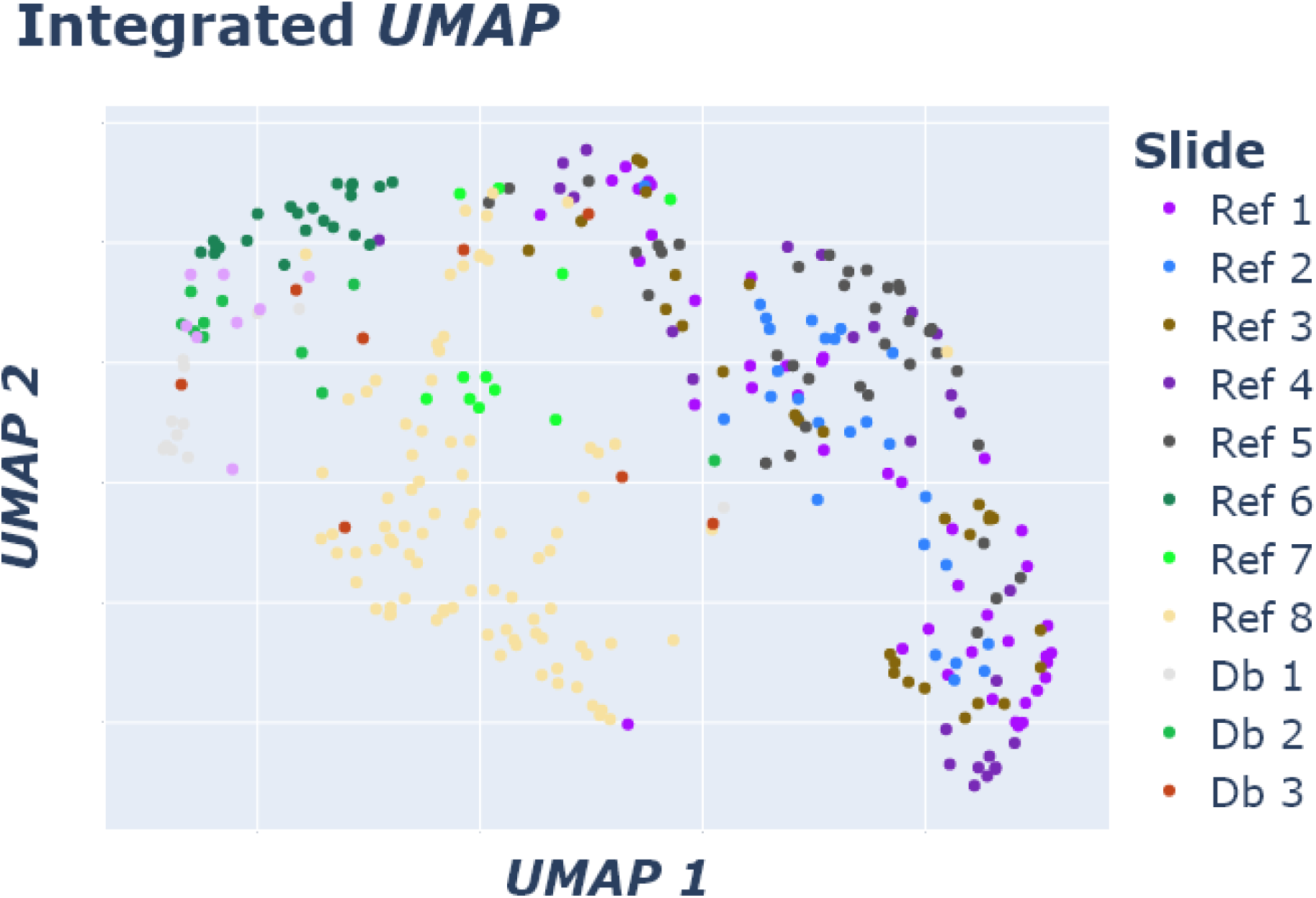
UMAP of morphometric and cell composition features from reference and diabetic slides. Each dot indicates a glomerulus.

**Supp. Fig. S6.**
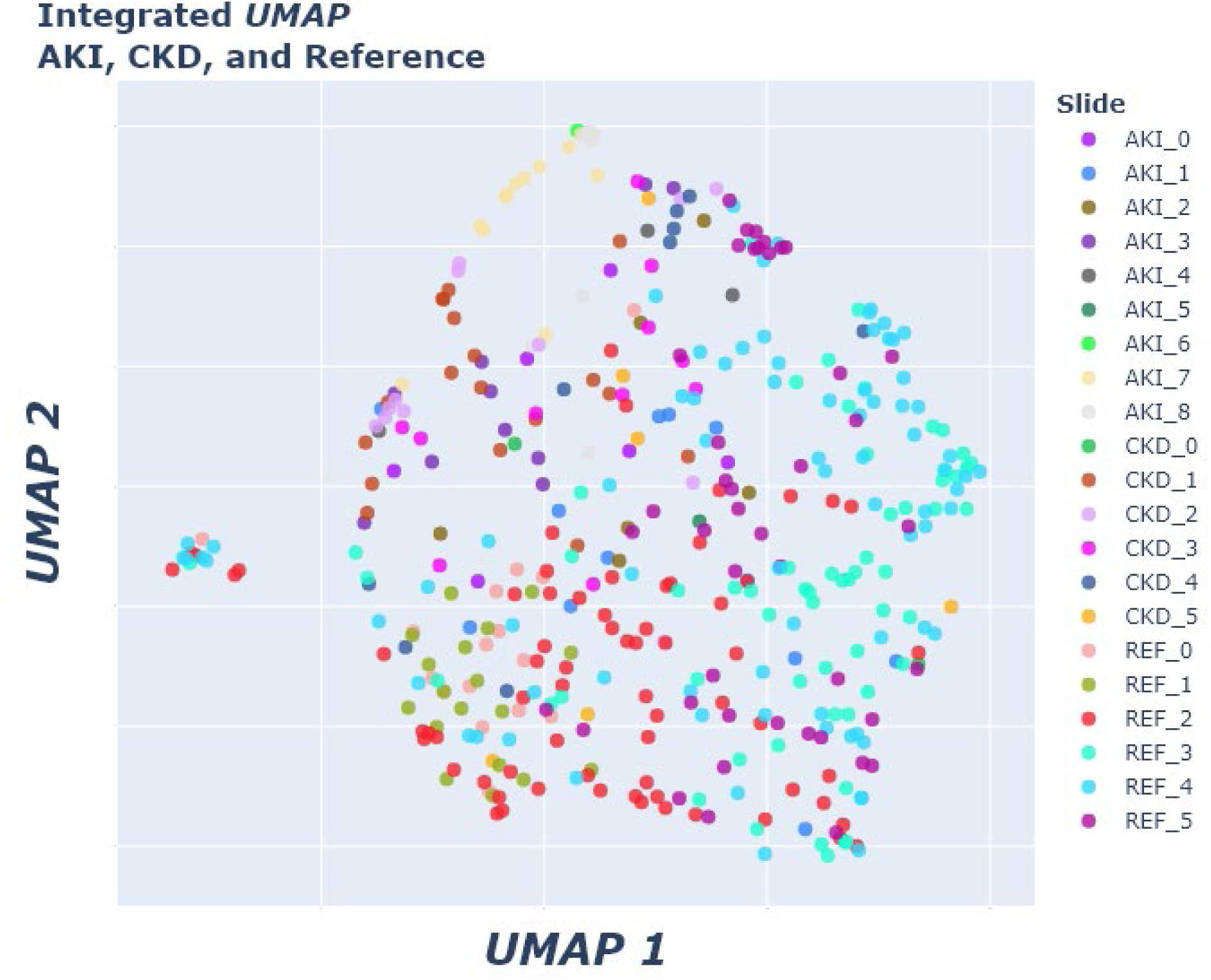
UMAP of morphometric features and cell composition from AKI, CKD, and reference slides. Each dot indicates a glomerulus.

### Use Case in Organs other than Kidney

To demonstrate the full utility of FUSION for other organs, one can use spatial data available for various reference organs from the HuBMAP portal. To increase the interoperability of our tool and better serve the HuBMAP community, FUSION is designed to accept HuBMAP processed datasets (.*h5ad* files) as well as ome-tiff formatted WSIs. One such example is shown for reference uterus tissue sections in ***Supp.*** Fig. 7. Users can select transcripts, whether that is the top-*k* highly variable transcripts, a list of named transcripts, or the top-*k* most prevalent transcripts. To gain further biological insights of these transcripts while navigating them spatially over tissue morphometry, FUSION leverages the HRA.^9^ The HRA includes information on molecular biomarkers, and their associated hierarchy of FTUs and organs in the human body. By integrating HRA’s publicly available API tools, FUSION can function as spatial guidebook for hypothesis generation for processed datasets imported from the HuBMAP portal.

**Supp. Fig. S7.**
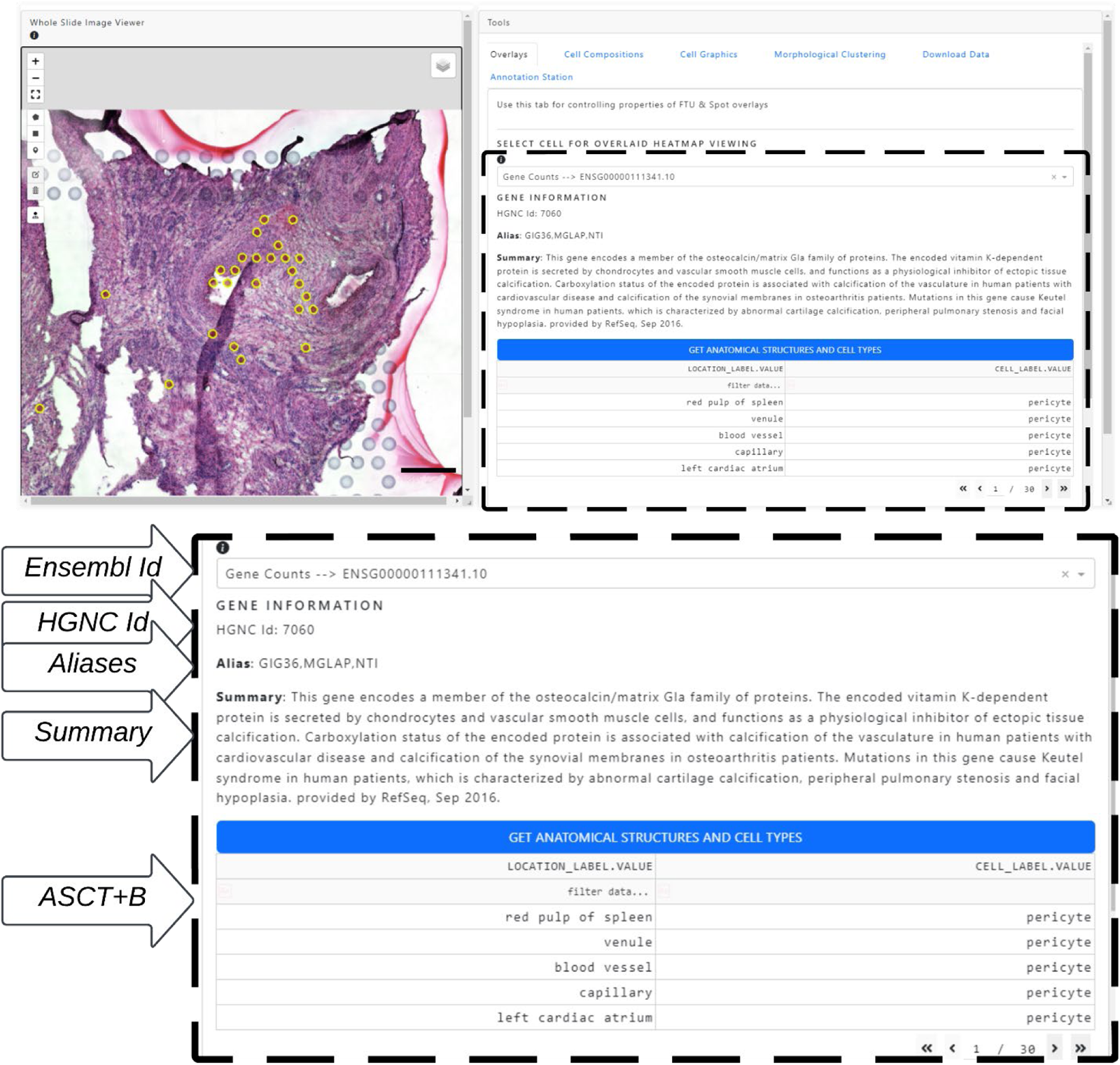
Biological queries using FUSION. Example usage of *10X Visium* data from a uterus section processed by the HuBMAP consortium and downloaded from their data portal. Enclosed section underneath heatmap overlay selection corresponds to information gathered pertaining to a particular gene along with a table with anatomical structures and cell types in the HRA that include that gene.^9^ Scalebar indicates 400 µm.

